# Combining genotypes and T cell receptor distributions to infer genetic loci determining V(D)J recombination probabilities

**DOI:** 10.1101/2021.09.17.460747

**Authors:** Magdalena L Russell, Aisha Souquette, David M Levine, E Kaitlynn Allen, Guillermina Kuan, Noah Simon, Angel Balmaseda, Aubree Gordon, Paul G Thomas, Frederick A Matsen, Philip Bradley

**Affiliations:** Computational Biology Program, Fred Hutchinson Cancer Research Center, Seattle, United States; Molecular and Cellular Biology Program, University of Washington, Seattle, United States; Department of Immunology, St. Jude Children’s Research Hospital, Memphis, United States; Department of Microbiology, Immunology, and Biochemistry, University of Tennessee Health Science Center, Memphis, United States; Department of Biostatistics, University of Washington, Seattle, United States; Centro Nacional de Diagnóstico y Referencia, Ministry of Health, Managua, Nicaragua; Sustainable Sciences Institute, Managua, Nicaragua; Department of Epidemiology, University of Michigan, Ann Arbor, United States; Institute for Protein Design, Department of Biochemistry, University of Washington, Seattle, United States

## Abstract

Every T cell receptor (TCR) repertoire is shaped by a complex probabilistic tangle of genetically determined biases and immune exposures. T cells combine a random V(D)J recombination process with a selection process to generate highly diverse and functional TCRs. The extent to which an individual’s genetic background is associated with their resulting TCR repertoire diversity has yet to be fully explored. Using a previously published repertoire sequencing dataset paired with high-resolution genome-wide genotyping from a large human cohort, we infer specific genetic loci associated with V(D)J recombination probabilities using genome-wide association inference. We show that V(D)J gene usage profiles are associated with variation in the *TCRB* locus and, specifically for the functional TCR repertoire, variation in the major histocompatibility complex locus. Further, we identify specific variations in the genes encoding the Artemis protein and the TdT protein to be associated with biasing junctional nucleotide deletion and N-insertion, respectively. These results refine our understanding of genetically-determined TCR repertoire biases by confirming and extending previous studies on the genetic determinants of V(D)J gene usage and providing the first examples of *trans* genetic variants which are associated with modifying junctional diversity. Together, these insights lay the groundwork for further explorations into how immune responses vary between individuals.

## Introduction

Receptor proteins on the surfaces of T cells are an essential component of the cell-mediated adaptive immune response in humans. Cells throughout the body regularly present protein fragments, known as antigens, on cell-surface molecules called major histocompatibility complex (MHC). Each T cell expresses a randomly-generated T cell receptor (TCR) which can bind the MHC- bound peptide and, if necessary, initiate an immune response. As part of this immune response, a T cell will proliferate and subsequent clones of that T cell will inherit the same antigen-specific TCR. Over time, the collection of all TCRs in an individual (the TCR repertoire) will dynamically summarize their previous immune exposures (***Woodsworth et al., 2013***).

To appropriately defend against a wide array of foreign pathogens, each individual has a highly diverse TCR repertoire. To generate diverse and functional TCRs, T cells combine a random generation process called V(D)J recombination with a selection process for proper expression and MHC recognition. Each TCR is composed of an *α* and a *β* protein chain which are both generated through V(D)J recombination. In the *β* chain, the recombination process proceeds by randomly choosing from a pool of V-gene, D-gene, and J-gene segments of the germline T cell receptor beta (*TCRB*) locus over a series of steps. First, the intervening chromosomal DNA between a randomly chosen D- and J-gene is removed to form a hairpin loop at the end of each gene (***Gellert, 1994**; **Fugmann et al., 2000**; **Schatz and Swanson, 2011***). Next, these hairpin loops are nicked open, often asymmetrically, by the Artemis-DNA-PKcs protein complex to create overhangs at the ends of each gene (***Weigert et al., 1978**; **Moshous et al., 2001**; **Ma et al., 2002**; **Lu et al., 2007**; **Zhao et al., 2020***). Depending on the location of the nick, the single-stranded overhang can contain short inverted repeats of gene terminal sequence known as P-nucleotides (***Nadel and Feeney, 1995**; **Gauss and Lieber, 1996**; **Nadel and Feeney, 1997**; **Jackson et al., 2004***). From here, nucleotides may be deleted from the gene ends through an incompletely-understood mechanism suggested to involve Artemis (***Feeney et al., 1994**; **Nadel and Feeney, 1995**, **1997**; **Jackson et al., 2004**; **Gu et al., 2010**; **Zhao et al., 2020***). This nucleotide trimming can remove traces of P-nucleotides (***Gauss and Lieber, 1996**; **Srivastava and Robins, 2012***). Next, non-templated nucleotides, known as N- insertions, can be added between the gene segments by the enzyme terminal deoxynucleotidyl transferase (TdT) (***Kallenbach et al., 1992**; **Gilfillan et al., 1993**; **Komori et al., 1993***). Once the nucleotide addition and deletion steps are completed, the gene segments are ligated together. The process is then repeated between this D-J junction and a random V-gene segment to generate a complete TCR*β* protein chain. After the *β* chain has been generated, a similar *α* chain recombination proceeds, although without a D-gene, to complete the TCR. Following the generation process, each completed TCR undergoes a selection process in the thymus to limit autoreactivity and ensure its ability to correctly bind peptide antigens presented on a specific MHC molecule (***Goldrath and Bevan, 1999**; **Thomas and Crawford, 2019***).

TCR repertoires vary between individuals and are a complicated tangle of genetically determined biases and immune exposures. Disentangling these factors is essential for understanding how our diverse repertoires support a powerful immune response. Previous efforts to unravel the genetic and environmental determinants governing TCR repertoire diversity have been highly impactful despite lacking high-throughput TCR repertoire sequencing data (***Sharon et al., 2016**; **Gao et al., 2019***) and/or high-resolution genotype data (***Rubelt et al., 2016**; **Emerson et al., 2017**; **Gao et al., 2019**; **Krishna et al., 2020***). For example, it has been shown that variation in the MHC locus biases TCR V(D)J gene usage (***Sharon et al., 2016**; **Gao et al., 2019***) and has been associated with clusters of shared receptors in response to Epstein-Barr virus epitope (***DeWitt et al., 2018***). Other studies have reported biases in V(D)J gene usage (***Zvyagin et al., 2014**; **Qi et al., 2016**; **Rubelt et al., 2016**; **Pogorelyy et al., 2018**; **Tanno et al., 2020**; **Fischer et al., 2021***), N-insertion lengths (***Rubelt et al., 2016***), and repertoire similarity in response to acute infection (***Qi et al., 2016; Pogorelyy et al., 2018***) for monozygotic twins. While this work clearly illustrates that genetic similarity implies TCR repertoire similarity, the extent to which specific variations are associated with V(D)J recombination probabilities has not been fully explored.

In this paper, we directly address the question of how an individual’s genetic background influences their V(D)J recombination probabilities using large human discovery and validation cohorts for which both TCR immunosequencing data (***Emerson et al., 2017**; **DeWitt et al., 2018***) and genotyping data (*Martin et al., 2020*) are available. With the goal of identifying statistically significant associations between single nucleotide polymorphisms (SNPs) and TCR repertoire features of interest using these novel, paired datasets, we treat analysis as a genome-wide association (GWAS) inference with many outcomes. Our results suggest that MHC and *TCRB* loci variations have an important role in determining the V(D)J gene usage profiles of each individual’s repertoire. At the junctions, we demonstrate that variations in the genes encoding the Artemis protein and the TdT protein are associated with biasing V- and J-gene nucleotide deletion and V-D and D-J-junction N-insertion, respectively.

## Results

### Discovery cohort data description

We worked with paired SNP array and TCR*β*-immunosequencing data representing 398 individuals and over 35 million SNPs and/or indels. TCR sequences can be separated into those that code for a complete, full-length peptide sequence (which we will call “productive” rearrangements) and “non-productive” rearrangements that do not. Non-productive sequences can arise during TCR generation steps if the V- and J-genes are shifted into different reading frames or a premature stop codon is introduced in the junction region. A non-productive rearrangement can be sequenced as part of the repertoire when a recombination event on one of a T cell’s two chromosomes fails to create a functional receptor, followed by a successful recombination event on the other chromosome. Because these non-productive sequences do not generate proteins that participate in the T cell selection process within the thymus, they should not be subject to functional selection (***Robins et al., 2010***). As such, their recombination statistics should reflect only the V(D)J recombination generation process which occurs before the stage of thymic selection.

In the data cohort of 398 individuals, an average of 235,054 unique TCR*β*-chain nucleotide sequences were sequenced per individual. Within each individual repertoire, roughly 18% of sequences were classified as “non-productive.” Thus, we can analyze the productive and non-productive sequences separately to distinguish between TCR generation and selection effects within each TCR repertoire. Specifically, we inferred the associations between genome-wide variation and V(D)J gene usage of each V-, D-, and J-gene, the extent of TCR nucleotide trimming, the number of TCR N-insertions, and the fraction of non-gene-trimmed TCRs containing P-nucleotides for both productive and non-productive sequences (***Table 1***).

**Table 1.** We inferred the associations between genome-wide variation and many different TCR repertoire features for productive and non-productive TCR sequences, separately. For each TCR repertoire feature, we considered the significance of associations using a Bonferroni-corrected threshold established to correct for each TCR feature subtype, the two TCR productivity types, and the total number of SNPs tested (described in detail in Methods).

| Repertoire feature<br>(significance threshold) | Model type | Feature subtype | Productivity | Significant association |
| --- | --- | --- | --- | --- |
| **V(D)J gene usage**<br>($4.72 \times 10^{-11}$) | simple | Each of 66 V-genes | Productive | Yes |
|  |  |  | Non-productive | Yes |
|  |  | Each of 2 D-genes | Productive | Yes |
|  |  |  | Non-productive | Yes |
|  |  | Each of 14 J-genes | Productive | Yes |
|  |  |  | Non-productive | Yes |
| **Amount of nucleotide trimming**<br>($9.68 \times 10^{-10}$) | gene-conditioned | V-gene trimming | Productive | Yes |
|  |  |  | Non-productive | Yes |
|  |  | 5' end D-gene trimming | Productive | No |
|  |  |  | Non-productive | No |
|  |  | 3' end D-gene trimming | Productive | No |
|  |  |  | Non-productive | No |
|  |  | J-gene trimming | Productive | Yes |
|  |  |  | Non-productive | Yes |
| **Number of N-insertions**<br>($1.94 \times 10^{-9}$) | simple | V-D-gene N-insertions | Productive | No |
|  |  |  | Non-productive | Yes |
|  |  | D-J-gene N-insertions | Productive | No |
|  |  |  | Non-productive | Yes |

### *TCRB* and MHC locus variation is associated with V-, D-, and J-gene usage frequency

To quantify the effect of SNPs on the expression of various V-, D-, and J-genes during V(D)J recombination, we designed a fixed effects model to assess the relationship between SNP genotype and gene frequency across all individuals. We fit this “simple model” for each different V-, D-, and J-gene in our paired dataset.

Because of the potential for population-substructure-related effects to inflate associations between each SNP and gene usage frequency, we incorporated ancestry-informative principal components (***Conomos et al., 2015***) based on the SNP genotypes for a subset of representative subjects as covariates in each model (see Methods for details). Diagnostic statistics show that this bias correction is sufficient (***Figure 1-source data 2***).

With these methods, we consider the significance of associations at a Bonferroni-corrected whole-genome P-value significance threshold of 4.72 × 10^−11^ (see Methods). Using this conservative threshold, we identified 9,957 significant associations between the frequency of various V-, D-, and J-genes and the genotype of SNPs genome-wide (***Figure 1*** and ***Figure 1-source data 1***). Of these significant associations, 7,678 were located within the *TCRB* locus for both productive and non-productive TCRs. The *TCRB* gene locus encodes the variable V-, D-, and J-gene segments which are recombined during V(D)J recombination. In our dataset, there are 66 V-genes, 2 D-genes, and 14 J-genes uniquely expressed. As we would expect, we find that the expression of many of these genes is associated with variation in the *TCRB* locus (***Figure 2***). Variation in the *TCRB* locus is most significantly associated with expression of the gene *TRBV28* within both productive and non-productive TCR*β* chains.

**Figure 1.**
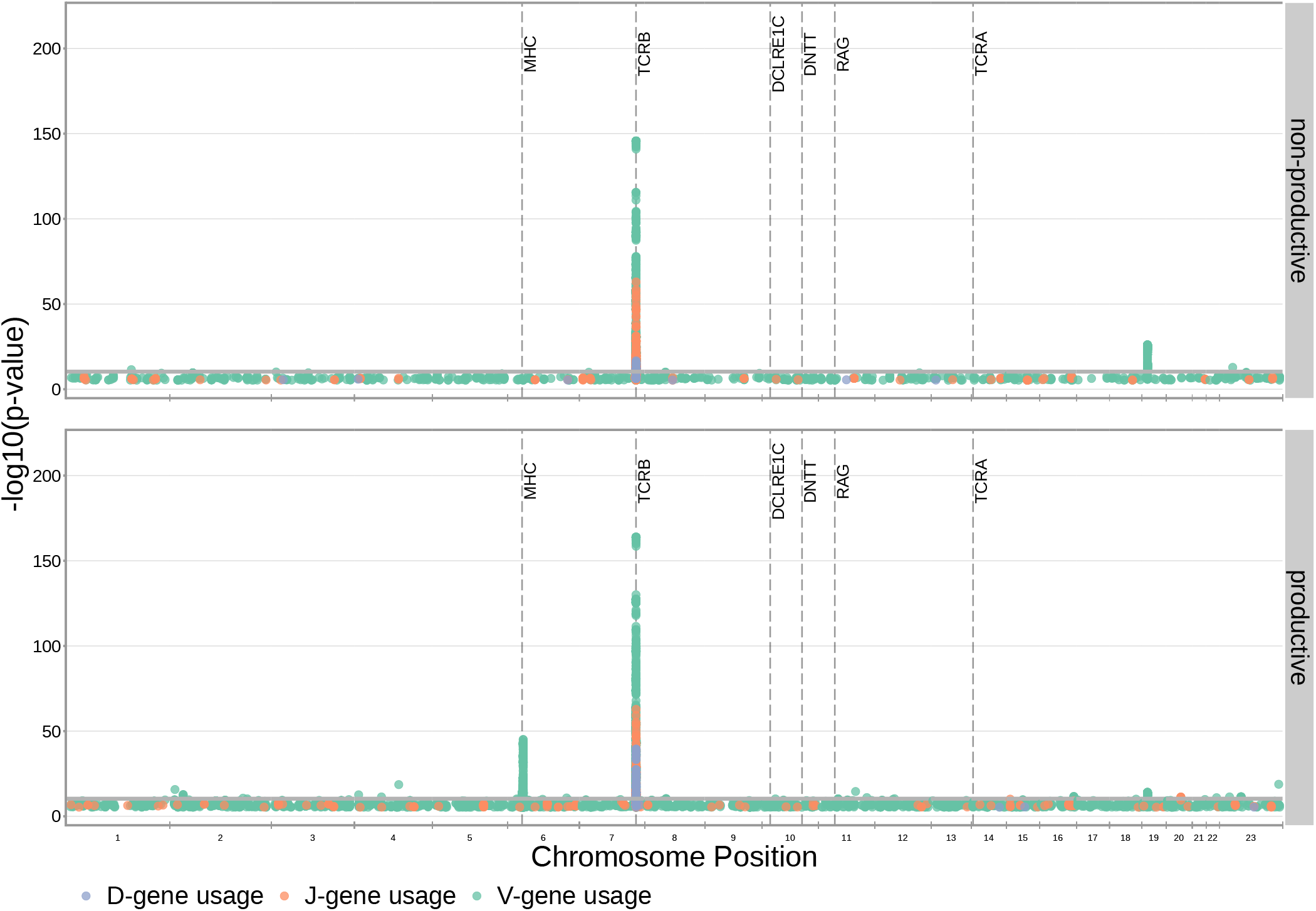
Many strong associations are present between V-, D-, and J-gene usage frequency and various SNPs genome-wide for both productive and non-productive TCRs. The most significant SNP associations for the frequency of each of the 66 V-genes, 2 D-genes, and 14J-genes are located within the *TCRB* and MHC loci. Associations are colored by gene-type instead of by gene identity for simplicity. Only SNP associations whose *P* < 5 × 10^−6^are shown here. The gray horizontal line corresponds to a Bonferroni-corrected P-value significance threshold of 4.72 ×10^−11^.

**Figure 2.**
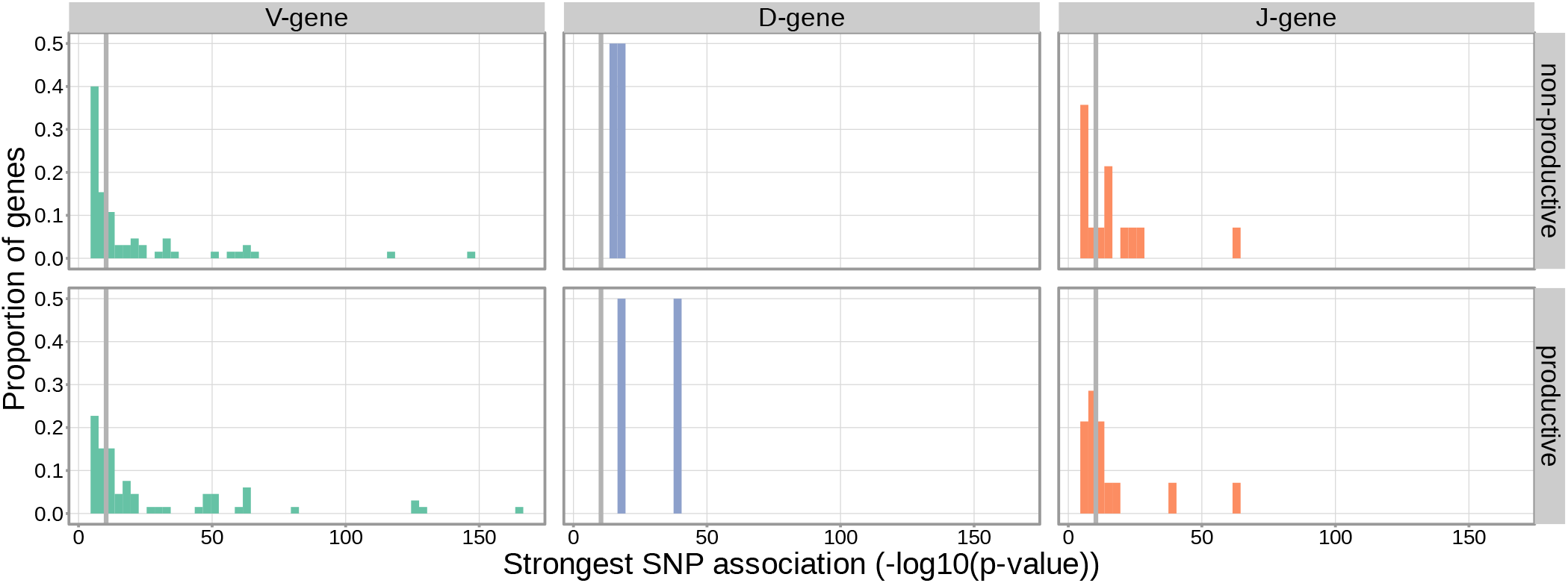
Gene-usage frequency of many V-gene, D-gene, and J-gene segments is significantly associated with variation in the *TCRB* locus. The P-value of the strongest *TCRB* SNP, gene-usage association for each different V-gene, D-gene, and J-gene segment is given on the X-axis. The proportion of gene segments within each gene type is given on the Y-axis. The gray vertical lines correspond to a whole-genome-level Bonferroni-corrected P-value significance threshold of 4.72 × 10^−11^.

Beyond the *TCRB* locus, we also identified 1,242 significant SNP associations within the major histocompatibility complex (MHC) locus. MHC proteins act by presenting self and foreign peptides to TCRs for inspection. Because of this important role in the functionality of T cells, the TCR-MHC interaction is important for thymic selection. We observe the expression of 12.1% of V-genes for productive TCRs to be associated with variation in the MHC locus. This associated MHC locus variation is located within sequences which code for canonical, peptide-presenting MHC proteins. For example, the eight most significantly associated SNPs were located within the *HLA-DRB1* gene within the MHC locus. These top SNPs were all associated with the expression of the gene *TRBV10-3* within productive TCRs. As expected, the expression of V-genes for non-productive TCRs is not associated with variation in the MHC locus. Likewise, the expression of D- and J-genes for both productive and non-productive TCRs is not associated with variation in the MHC locus. These results refine and extend associations found in previous work (***Sharon et al., 2016**; **Gao et al., 2019***).

We observed just one other long-range association region, in addition to the MHC locus, located in proximity to the *ZNF443* and *ZNF709* loci on chromosome 19. Both of these zinc finger proteins contain KRAB-domains and, thus, likely act as transcriptional repressors (***Witzgall et al., 1994***). In this region, we observe 1,004 significant SNP associations for the expression of the V-genes *TRBV24-1*and *TRBV24/OR9-2*. Of these 1,004 SNP associations, 604 were associations for V-gene expression in non-productive TCRs and 400 were associations for V-gene expression in productive TCRs. Because these associations are strongest for non-productive TCRs, this chromosome 19 variation likely influences gene usage during TCR generation steps, as opposed to selection. Variation in proximity to the *ZNF443* and *ZNF709* loci may alter the resulting zinc finger proteins and lead to differential transcriptional repression of a site near *TRBV24*. Because the transcription of unrearranged gene segments influences their recombination potential (***Oltz, 2001***), this difference in repression could subsequently change the usage frequency of the *TRBV24* gene.

### *DCLRE1C* locus variation is associated with the extent of V-, D-, and J-gene trimming

We hypothesized that SNPs across the genome, particularly those within V(D)J-recombination-associated genes, may influence the extent of TCR nucleotide trimming at V(D)J *TCRB* gene junctions. It has been previously observed that the extent of trimming varies by V(D)J *TCRB* gene choice (***Figure 3-Figure Supplement 4***) (***Nadel and Feeney, 1995**, **1997**; **Jackson et al., 2004**; **Murugan et al., 2012***). In other words, two different V-genes (*TRBV19* and *TRBV20-1*, for example) will on average be trimmed to different extents due, in part, to differences in their terminal nucleotide sequences (and the same is true for D- and J-genes). Thus, to quantify the effect of SNPs on the extent of V-, D-, and J-gene trimming during V(D)J recombination, without confounding the extent of trimming with *TCRB* gene choice, we designed a linear fixed effects model to measure the correlation between each SNP and the number of nucleotide deletions, while conditioning out the effect mediated by gene choice. We fit this “gene-conditioned model” for each of the four trimming types (V-gene trimming, 5′ end D-gene trimming, 3′ end D-gene trimming, and J-gene trimming) on our paired data set. We performed the analysis, as above, incorporating ancestry-informative principal components in each model (detailed in Methods). Diagnostic statistics show that this correction for population-substructure-related biases is sufficient (***Figure 3-source data 2***). Here, we considered the significance of associations at a Bonferroni-corrected whole-genome P-value significance threshold of 9.68 × 10^−10^ (see Methods).

With these methods, we identified 317 significant SNP associations with the extent of nucleotide trimming for various trimming types (***Figure 3*** and ***Figure 4-source data 1***). We found 66 highly significant associations between V- and J-gene trimming and SNPs within the *DCLRE1C* gene locus for both productive and non-productive TCRs when considered in the whole-genome context. The *DCLRE1C* gene encodes the Artemis protein, an endonuclease responsible for cutting the hairpin intermediate prior to nucleotide trimming and insertion during V(D)J recombination. Many of the SNPs responsible for these 66 significant associations within the *DCLRE1C* locus were shared between trimming and productivity types (***Figure 4***). The most significantly-associated SNP (rs41298872) within this locus had a P-value of 3.18 × 10^−37^ for J-gene trimming of non-productive TCRs (***Figure 3-Figure Supplement 1***). This SNP was also significantly-associated with J-gene trimming of productive (*P* = 1.99 × 10^−29^) TCRs and V-gene trimming of productive (*P* = 6.23 × 10^−23^) and non-productive (*P* = 2.81 × 10^−21^) TCRs. We performed a conditional analysis to identify potential independent secondary signals by including this SNP as an additional covariate within the model. This analysis revealed a second, independent SNP signal (rs35441642) within the *DCLRE1C* locus forJ-gene trimming of non-productive TCRs (***Figure 4-source data 2***). None of the other nucleotide trimming type, productivity status combinations had significant evidence for secondary independent signals.

**Figure 3.**
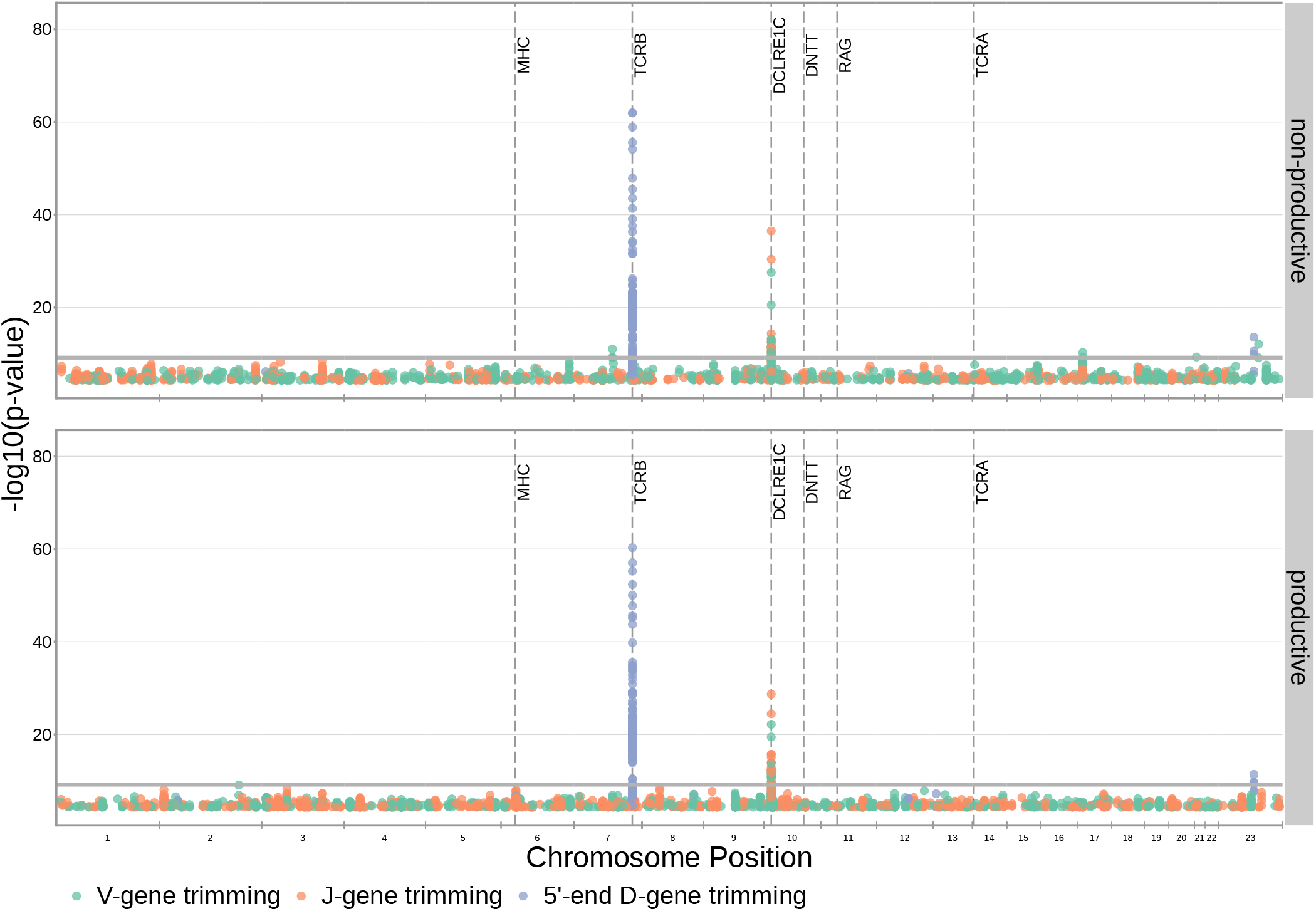
SNP associations for all four trimming types reveal the most significant associations to be located within the *TCRB* and *DCLRE1C* loci for 5′ D-gene trimming and V-gene trimming, respectively, when conditioning out effects mediated by gene choice when calculating the strength of association. Only SNP associations whose *P* < 5 × 10^−5^ are shown here. The gray horizontal line corresponds to a Bonferroni-corrected P-value significance threshold of 9.68 × 10^−10^.

**Figure 4.**
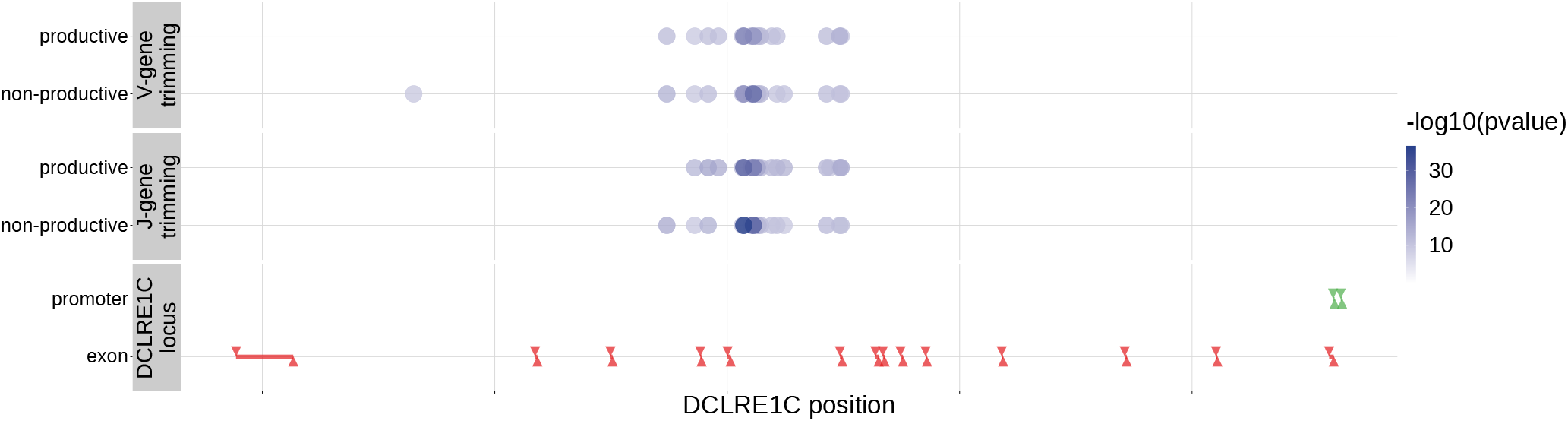
Within the *DCLRE1C* locus, 93.8% of these significantly associated SNPs were located within introns. Additionally, many of these significant SNP associations overlapped between trimming types. Downward arrows represent promoter/exon starting positions and upward arrows represent promoter/exon ending positions.

Our procedure also identified many highly significant associations between 5’ end D-gene trimming and SNPs within the *TCRB* gene locus, however these appear to result from correlations between SNP genotype and *TRBD2* allele genotype (***Figure 3-Figure Supplement 1***). If we correct for *TRBD2* allele genotype in our model formulation (see Methods), we no longer observe these associations between SNPs within the *TCRB* gene locus and the extent of 5′ end D-gene trimming (***Figure 3-Figure Supplement 2***). *TRBD2* allele genotype could be acting as a confounding variable due to linked local genetic variation which influences nucleotide trimming and/or D-gene assignment ambiguity variation as a function of *TRBD2* allele genotype. To explore the extent of possible D-gene assignment ambiguity variation, we restricted our analysis to TCRs which contain *TRBJ1* genes (and consequently contain *TRBD1* due to topological constraints during V(D)J recombination (***Robins et al., 2010**; **Murphy and Weaver, 2016***)). With this approach, we also no longer observe associations between SNPs within the *TCRB* gene locus and the extent of 5′ end D-gene trimming, and additionally, we do observe significant associations between SNPs within the *DCLRE1C* locus and 5′ and 3’ end D-gene trimming which were not observed in the original genome-wide analysis (***Figure 3-Figure Supplement 3***).

Our fixed effects model formulation for these inferences is important: if we don’t condition on gene choice then additional, and presumably spurious, associations arise. Indeed, when implementing the “simple model” designed to quantify the association between the four trimming types and genome-wide SNP genotypes, without conditioning out the effect mediated by gene choice, we observe additional associations between SNPs within the MHC locus and V-gene trimming of productive TCRs and between SNPs within the *TCRB* locus and V-gene and 3’ end D-gene trimming of, again, productive TCRs (***Figure 3-Figure Supplement 5***). This is perhaps not surprising, as we noted earlier that variations in the MHC and *TCRB* loci are associated with gene usage frequencies in productive TCRs (***Figure 1***), and different genes have different trimming distributions (determined in part by the nucleotide sequences at their termini).

Because P-nucleotides can be present at V(D)J junctions in the absence of nucleotide trimming (***Murphy and Weaver, 2016***), we hypothesized that similar *DCLRE1C* locus variation may also be associated with P-addition. Interestingly, we did not identify any strong associations between SNPs within the *DCLRE1C* locus and the fraction of non-gene-trimmed TCRs containing P-nucleotides when implementing our “gene-conditioned model”, despite the known role of the Artemis protein in functioning as an endonuclease responsible for cutting the hairpin intermediate, and thus, potentially creating P-nucleotides during V(D)J recombination (***Figure 3-Figure Supplement 6***). We observe similar results when quantifying the effect of genome-wide SNPs on the number of V-, D-, and J-gene P-nucleotides per TCR (***Figure 3-Figure Supplement 7***).

### *DNTT* locus variation is associated with the number of V-D and D-J N-insertions

Unlike V-, D-, or J-gene nucleotide trimming length, the number of nucleotide N-insertions between V-D and D-J genes does not vary substantially with V(D)J *TCRB* gene choice (***Figure 5-Figure Supplement 1***) (***Murugan et al., 2012***). Thus, to infer the association between SNPs and the number of nucleotide N-insertions, we implemented a “simple model”, without conditioning out any effect mediated by gene choice. Again, because of the potential for population-substructure-related effects to inflate associations between each SNP and the number of N-insertions, we incorporated ancestry-informative principal components as covariates in each model (detailed in Methods). Diagnostic statistics show that this bias correction is sufficient (***Figure 5-source data 3***).

With these methods, we identified three associations between SNPs and the number of nucleotide N-insertions using a Bonferroni-corrected whole-genome P-value significance threshold of 1.94 × 10^−9^ (see Methods) (***Figure 5*** and ***Figure 5-source data 1***). Two SNPs within the *DNTT* gene locus (rs2273892 and rs12569756) were responsible for these associations. The *DNTT* gene encodes the terminal deoxynucleotidyl transferase (TdT) protein which is a specialized DNA polymerase responsible for adding non-templated (N) nucleotides to coding junctions during V(D)J recombination. When we restrict our analysis to TCRs which contain *TRBJ1* genes and consequently eliminate potential D-gene assignment ambiguity, we continue to observe these *DNTT* associations (***Figure 5-Figure Supplement 2***).

**Figure 5.**
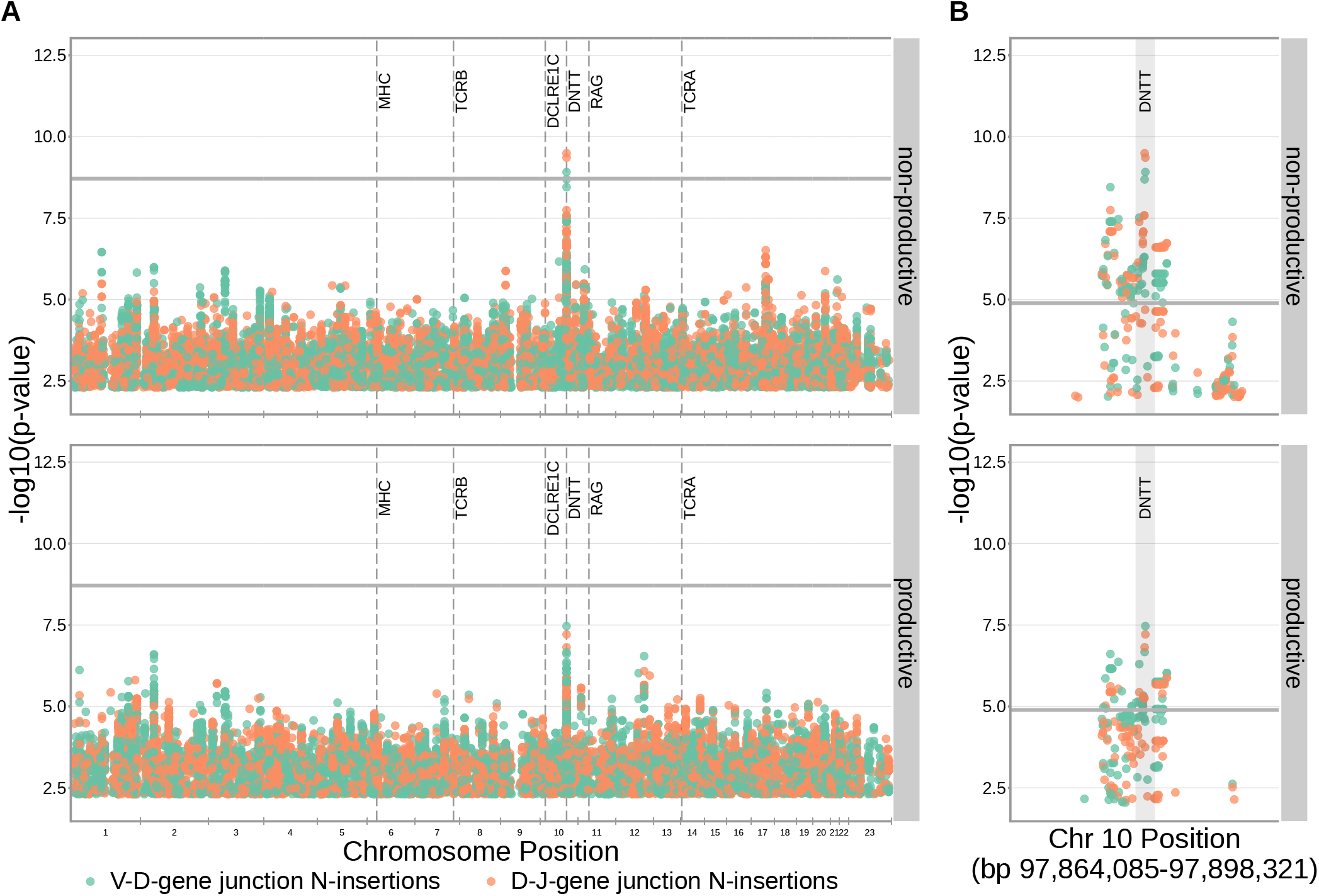
SNPs within the *DNTT* locus are associated with the extent of N-insertion. (**A**) There are three associations for SNPs within the *DNTT* locus which are significant when considered in the whole-genome context. The gray horizontal line corresponds to a whole-genome Bonferroni-corrected P-value significance threshold of 1.94 × 10^−9^. (**B**) Using a *DNTT* gene-level significance threshold, many more SNPs within the extended *DNTT* locus have significant associations for both N-insertion types. Here, the gray horizontal line corresponds to a gene-level Bonferroni-corrected P-value significance threshold of 1.28 × 10^−5^ (calculated using gene-level Bonferroni correction for the 977 SNPs within 200kb of the *DNTT* locus, see Methods). For both (A) and (B), only SNP associations whose *P* < 5 × 10^−3^ are shown.

Since the TdT protein has an important mechanistic role in the N-insertion process and because we already identified SNPs within the *DNTT* locus to be weakly associated with the number of N-insertions at V(D)J gene junctions, we wanted to explore the locus further. Restricting the analysis to the extended *DNTT* locus reduced the multiple testing burden such that 232 significant associations emerged (***Figure 5*** and ***Figure 5-source data 2***). Many of the SNPs responsible for these 232 significant associations within the extended *DNTT* locus were shared between insertion and productivity types (***Figure 6***). While most of these associations are likely the result of a single independent signal for each insertion and productivity type, we performed a conditional analysis to identify potential independent secondary signals. To do so, we included the most significant SNP within the *DNTT* locus for each insertion and productivity type as a covariate in the model. With this approach, we identified rs2273892 as the primary independent signal for D-J N-insertion of non-productive TCRs and rs12569756 as the primary independent signal for D-J N-insertion of productive TCRs and V-D N-insertion of productive and non-productive TCRs. However, these two SNPs are tightly linked and, thus, likely both represent the same, primary independent signal. This analysis did not reveal any significant evidence for secondary independent signals.

**Figure 6.**
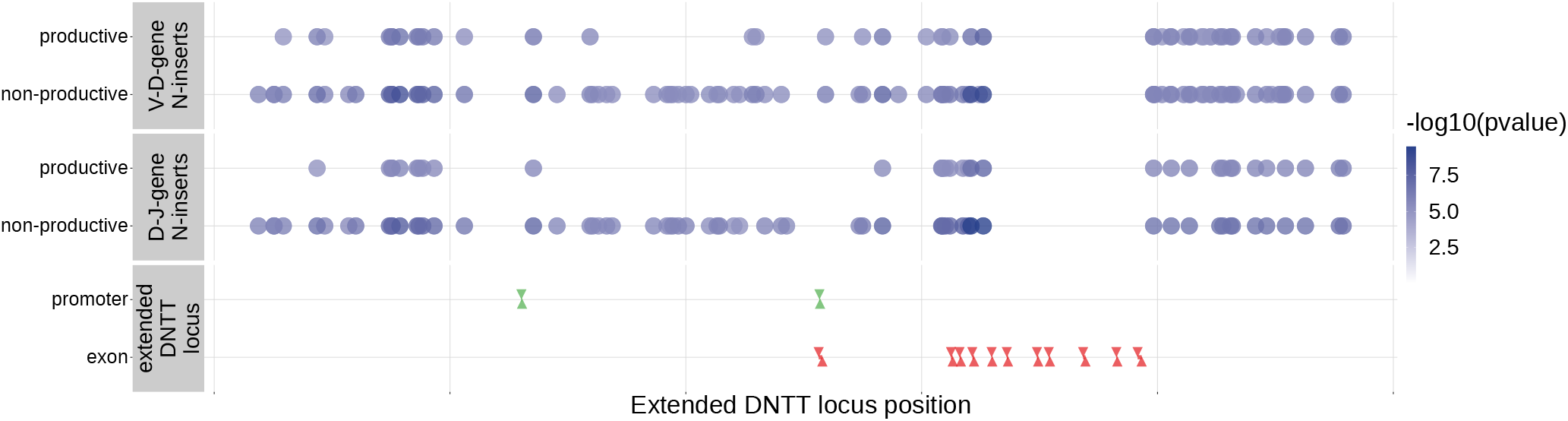
Within the *DNTT* locus, many of the significant SNP associations overlapped between N-insertion types when using *DNTT* gene-level Bonferroni-corrected P-value significance threshold of 1.28 × 10^−5^. Downward arrows represent promoter/exon starting positions and upward arrows represent promoter/exon ending positions.

We found that correcting for population-substructure-related effects was especially important in our primary genome-wide analysis, which led us to discover differences in the extent of N- insertion by ancestry-informative PCA cluster. Indeed, if we don’t incorporate correction terms for population-substructure-related biases in our model formulation, we observe many strongly significant associations, particularly within the *DNTT* locus. This hinted at important PCA-cluster level effects. When we look closely at the average number of N-insertions (combining the number of V-D and D-J N-insertions) across TCR repertoires by PCA cluster, we note that subjects from the “Asian”-associated PCA cluster have significantly fewer total N-insertions for productive (*P* = 0.006 without Bonferroni correction) and non-productive (*P* = 0.014 without Bonferroni correction) TCRs when compared to the population mean (using a one-sample t-test) (***Figure 7***). The total N-insertions for productive TCRs within the “Asian”-associated PCA cluster remain significantly different from the population mean after Bonferroni multiple testing correction (corrected *P* = 0.036). Furthermore, the “Asian”- and “Hispanic”-associated PCA clusters had significantly higher mean SNP minor allele frequencies for N-insertion associated SNPs within the extended *DNTT* region when compared to the mean population minor allele frequency (*P* = 7.32 × 10^−20^ for the “Asian”-associated PCA cluster and *P* = 1.17 × 10^−5^ for the “Hispanic”-associated PCA cluster using a one-sample t-test with Bonferroni multiple testing correction) (***Figure 8***).

**Figure 7.**
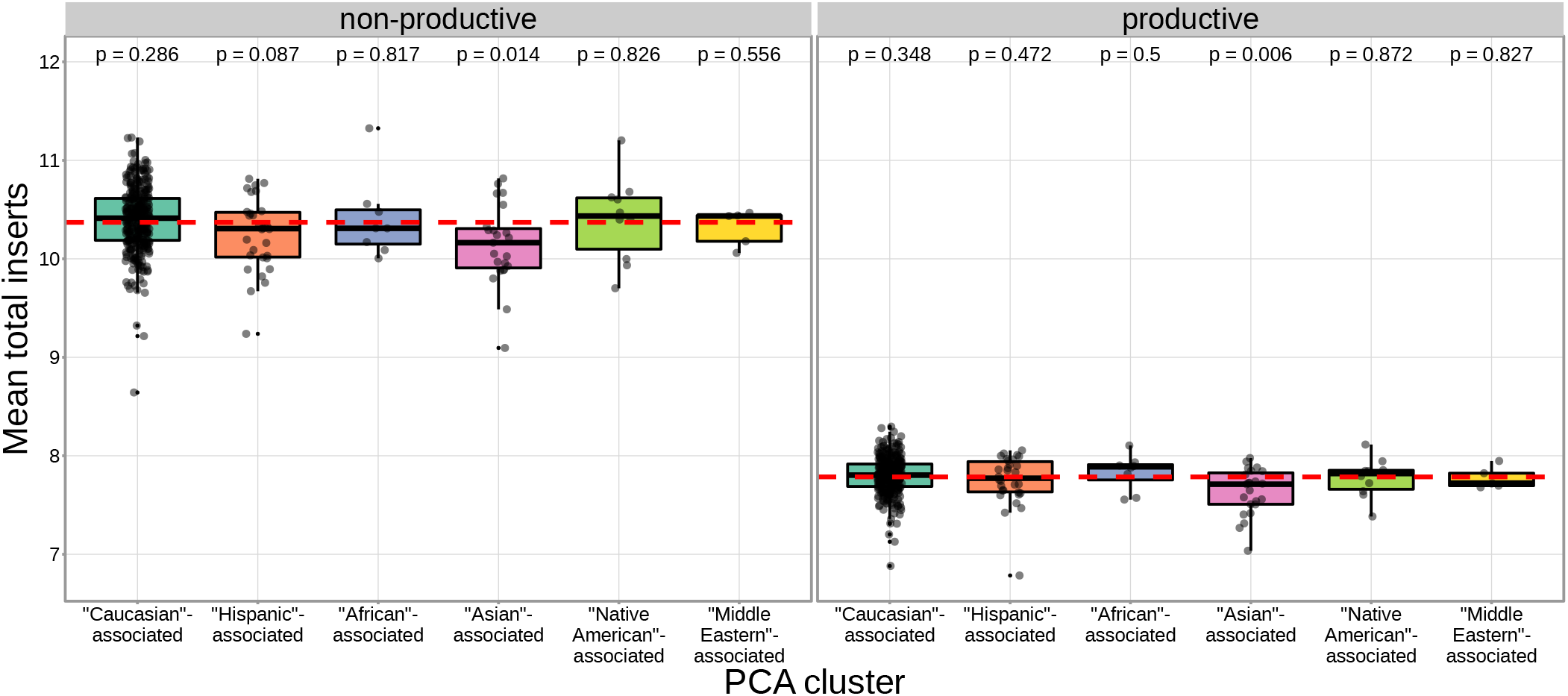
The TCR repertoires for subjects in the “Asian”-associated PCA-cluster contain fewer N-insertions for productive TCRs when compared to the population mean computed across all 666 subjects (dashed, red horizontal line). The P-values from a one-sample t-test (without Bonferroni multiple testing correction) for each PCA cluster compared to the population mean are reported at the top of the plot.

**Figure 8.**
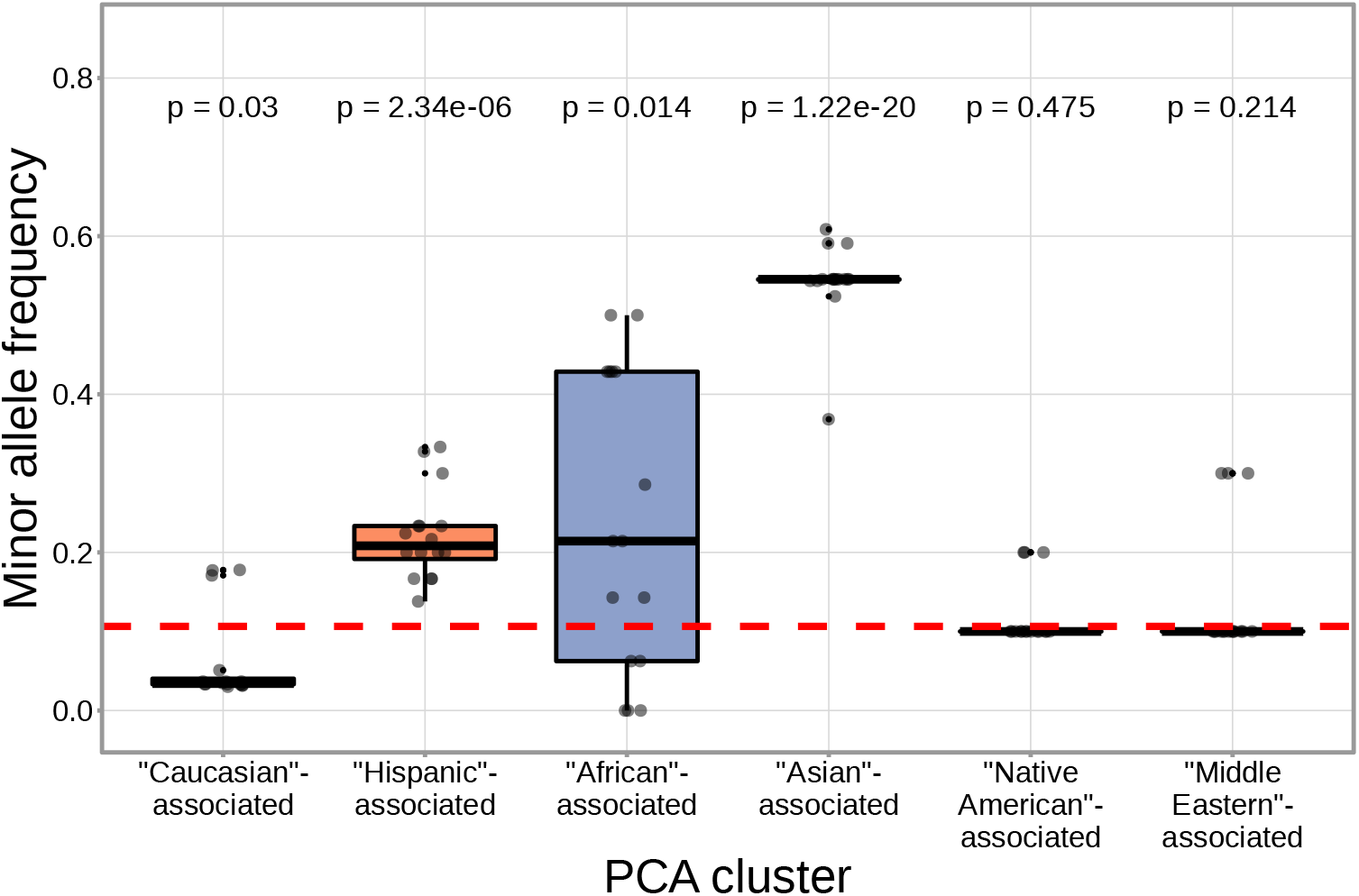
N-insertion associated SNPs within the *DNTT* region have a higher mean minor allele frequency within the “Asian”-associated PCA-cluster when compared to the population mean minor allele frequency computed across the 398 discovery cohort subjects (dashed, red horizontal line). The P-values from a one-sample t-test (without Bonferroni multiple testing correction) for each PCA cluster compared to the population mean are reported at the top of the plot. The population mean is dominated by subjects in the “Caucasian”-associated PCA cluster (***Figure 7-Figure Supplement 1***).

### Validation Analysis

To validate our results, we worked with paired ancestry-informative marker (AIM) SNP array and TCR*α*- and TCR*β*-immunosequencing data representing 94 individuals and 2 SNPs (which overlap with the discovery dataset) from an independent validation cohort (see Methods). In contrast to the discovery cohort, this cohort contains different demographics, shallower RNA-seq based TCR-sequencing, and a sparser set of SNPs. However, TCR-sequencing for both TCR*α* and TCR*β* chains is available.

We were able to validate a discovery-cohort significantly associated *DCLRE1C* SNP within this validation cohort. While none of the independent *DCLRE1C* SNPs from the discovery-cohort analysis overlapped with the validation cohort SNP set, a single, non-synonymous SNP (rs12768894, c.728A>G) within the *DCLRE1C* locus was present in both SNP sets. This SNP was one of the significant associations we observed for V-gene trimming (productive *P* = 2.16 × 10^−14^; non-productive *P* = 7.21 × 10^−14^) and J-gene trimming (productive *P* = 1.23 × 10^−11^; non-productive *P* = 6.62 × 10^−12^) of TCR*β* chains in the genome-wide discovery cohort analysis (***Table 2-Figure Supplement 1***). Using the same methods, we identified significant associations between this SNP and J-gene trimming of productive TCR*α* and TCR*β* chains and V-gene trimming of both productive and non-productive TCR*α* and TCR*β* chains within the validation cohort (***Table 2**, **Table 2-Figure Supplement 2***, and ***Table 2-Figure Supplement 3***). Associations between rs12768894 and both types of D-gene trimming of TCR*β* chains were not significant for either cohort.

**Table 2.** We inferred the associations between SNP genotype and TCR repertoire features for two SNPs overlapping between discovery-cohort and validation-cohort SNP sets. We considered the significance of the validation cohort associations at a Bonferroni-corrected SNP-level P-value significance threshold of 0.0042 for trimming and 0.0083 for N-insertion (see Methods). Validation cohort P-values are one-tailed. * discovery-cohort associations were only significant when considered at the *DNTT* -gene level significance threshold, not at the whole-genome significance threshold.

| SNP | TCR chain | Repertoire feature | Productivity type | Discovery cohort signif. association | Validation cohort signif. association |
| --- | --- | --- | --- | --- | --- |
| rs12768894 | TCRβ | V-gene trimming | Productive | Yes ($2.16 \times 10^{-14}$) | Yes ($7.17 \times 10^{-7}$) |
| | | | Non-productive | Yes ($7.21 \times 10^{-14}$) | Yes ($8.75 \times 10^{-6}$) |
| | | J-gene trimming | Productive | Yes ($1.23 \times 10^{-11}$) | Yes ($5.16 \times 10^{-10}$) |
| | | | Non-productive | Yes ($6.62 \times 10^{-12}$) | No ($4.18 \times 10^{-2}$) |
| | TCRα | V-gene trimming | Productive | N/A | Yes ($2.59 \times 10^{-5}$) |
| | | | Non-productive | N/A | Yes ($2.68 \times 10^{-7}$) |
| | | J-gene trimming | Productive | N/A | Yes ($6.29 \times 10^{-12}$) |
| | | | Non-productive | N/A | No ($9.99 \times 10^{-3}$) |
| rs3762093 | TCRβ | V-D N-insertion | Productive | Yes* ($1.37 \times 10^{-6}$) | No (0.153) |
| | | | Non-productive | Yes* ($1.50 \times 10^{-7}$) | No (0.059) |
| | | D-J N-insertion | Productive | Yes* ($9.43 \times 10^{-6}$) | No (0.137) |
| | | | Non-productive | Yes* ($1.94 \times 10^{-7}$) | No (0.067) |
|  | TCRα | V-J N-insertion | Productive | N/A | Yes (0.006) |
|  |  |  | Non-productive | N/A | No (0.031) |

We were unable to validate the most significantly associated *DNTT* SNPs due to lack of overlap between the SNP sets for the discovery and validation cohorts; a discovery-cohort weakly associated SNP (rs3762093) failed to reach statistical significance for all N-insertion types, but had the same direction of effect in the validation cohort as follows. Within the discovery cohort, rs3762093 genotype was weakly associated with the number of V-D N-insertions (productive *P* = 1.37 × 10^−6^; non-productive *P* = 1.50 × 10^−7^) and D-J N-insertions (productive *P* =9.43 × 10^−6^; non-productive *P* = 1.94 × 10^−7^) within TCR*β* chains (***Table 2-Figure Supplement 4***). Within the validation cohort, this SNP was significantly associated with the number of V-J N-insertions within productive TCR*α* chains (***Table 2*** and ***Table 2-Figure Supplement 6***). However, this SNP was not significantly associated with the number of V-D or D-J N-insertions within productive or non-productive TCR*β* chains or the number of V-J N-insertions within non-productive TCR*α* chains within the validation cohort (***Table 2**, **Table 2-Figure Supplement 5***, and ***Table 2-Figure Supplement 6***). Despite the lack of significance, we noted that the model coefficients for rs3762093 genotype were in the same direction (i.e., the minor allele was associated with fewer N-insertions) for all N-insertion and productivity types within TCR*β* chains for both cohorts. Further, while TCR*α* chain sequencing was not available for the discovery cohort, we observed stronger associations between rs3762093 genotype and the extent of N-insertion for both productivity types within TCR*α* chains compared to TCR*β* chains within the validation cohort. Perhaps with a larger validation cohort, significant associations would be present for all N-insertion types.

## Discussion

V(D)J recombination is a complex stochastic process that enables the generation of diverse TCR repertoires. Our results show that genetic variation in various V(D)J recombination genes has a key role in shaping the TCR repertoire through biasing V(D)J gene choice, nucleotide trimming, and N-insertion in a broad population sample. While we recognize that there may be a complicated entanglement between allelic variation and local *cis*-acting effects, we were primarily interested in identifying strong, *trans*-acting associations. By leveraging the unique pairing of TCR*β* chain immunosequencing and genome-wide genotype data, we have (1) confirmed and extended previous studies on the genetic determinants of TCR V-gene usage, (2) discovered associations between common genetic variants within the *DCLRE1C* and *DNTT* loci and V(D)J junctional trimming and N-insertions, respectively, (3) developed a method for quantifying the extent of the associations between genetic variations and junctional features, directly, without confounding gene choice effects, and (4) revealed differences in the extent of N-insertion by ancestry-informative PCA cluster.

We note an abundance of associations between variation in the *TCRB* locus and V(D)J gene usage biases for both productive and non-productive TCRs. Although previous reports have revealed similar patterns of association for productive TCRs (***Sharon et al., 2016**; **Gao et al., 2019***), our results refine and extend this result by quantifying the extent of *TCRB* locus variation on V(D)J gene usage for non-productive TCRs. This highlights that locus variation is associated with TCR generation-related gene usage biases, in addition to potential thymic selection biases for productive TCRs. These TCR generation-related gene usage biases likely reflect local gene regulation and/or recombination efficiency effects. For example, one of the SNPs most significantly associated with *TRBV28* expression (rs17213) is located within the recombination signal sequence at the 3′-end of the gene and, thus, could be involved directly in changing the recombination efficiency of *TRBV28*. Thus, different expression levels of various genes could be promoted by variation within non-coding regions such as promoters, 5’UTRs and leader sequences, introns, or recombination signal sequences. Polymorphisms within these regions have been suggested to influence V(D)J gene expression levels within B-cell receptor repertoires (***Mikocziova et al., 2021***). We also observed that variation in the MHC locus is associated with V-gene usage biases for productive TCRs, but not non-productive TCRs. These MHC locus associations are likely only observed for V-gene usage since the V-gene locus, exclusively, encodes the TCR regions (complementarity-determining regions 1 and 2) which directly contact MHC during peptide presentation (***Murphy and Weaver, 2016***). Previous work has suggested that the thymic selection of certain V-genes may be biased by germline-encoded TCR-MHC compatibilities in an MHC dependent manner (***Sharon et al., 2016**; **Gao et al., 2019***). Because of our observed distinction between associations present between MHC variation and V-gene usage in productive versus non-productive TCRs, our work supports this hypothesis.

We have identified, for the first time, specific genetic variants which are associated with modifying the extent of N-insertion and nucleotide trimming. While many previous studies have reported evidence of genetic influences on overall gene usage (***Zvyagin et al., 2014**; **Qi et al., 2016**; **Rubelt et al., 2016**; **Pogorelyy et al., 2018**; **Tanno et al., 2020**; **Fischer et al., 2021***) and repertoire similarity in response to acute infection (***Qi et al., 2016**; **Pogorelyy et al., 2018***), there have been few explorations into how heritable factors may bias TCR junctional features beyond reports of genetic similarity implying overall TCR repertoire similarity (***Krishna et al., 2020**; **Rubelt et al., 2016***). Here, we noted that variation in the gene encoding the Artemis protein (*DCLRE1C*) is associated with the extent of V- and J-gene nucleotide trimming for both productive and non-productive TCRs. These associations are strongest for non-productive TCRs suggesting a TCR generation-related repertoire bias. It is well established that the Artemis protein, in complex with DNA-PKcs, functions as an endonuclease responsible for cutting the hairpin intermediate, and thus, potentially creating P-nucleotides prior to nucleotide trimming during V(D)J recombination (***Weigert et al., 1978**; **Moshous et al., 2001**; **Ma et al., 2002**; **Lu et al., 2007***). The direct involvement of Artemis in the nucleotide trimming mechanism, however, has yet to be confirmed. It has been shown that the Artemis protein possesses single-strand-specific 5′ to 3′ exonuclease activity (***Ma et al., 2002**; **Li et al., 2014***) and, thus, may be properly positioned to trim nucleotides. A non-synonymous SNP within *DCLRE1C* (rs12768894, c.728A>G) was one of the significant associations we observed forV- and J-gene nucleotide trimming in both the primary cohort and the independent validation cohort. Perhaps this mutation, or other linked non-synonymous *DCLRE1C* variation that was not studied here, is directly involved in the trimming changes we observe. We did not observe strong associations between variation in the *DCLRE1C* locus and the number of P-nucleotides or the fraction of non-gene-trimmed TCRs containing P-nucleotides, despite the established mutually exclusive relationship between P-addition and nucleotide trimming (***Gauss and Lieber, 1996**; **Srivastava and Robins, 2012**; **Murphy and Weaver, 2016***). However, the absence of P-nucleotide associations at the *DCLRE1C* locus could be the result of restricting the analyses to the non-gene-trimmed repertoire subset. Perhaps with a larger dataset these associations would be present.

Further, we have identified associations between variation in the gene encoding the TdT protein (*DNTT*) and the number of N-insertions for both productive and non-productive TCRs. Because of the established, direct involvement of the TdT protein in the N-insertion mechanism, these *DNTT* locus variations could be influencing the function of the TdT protein. These significant associations were slightly stronger for non-productive TCRs perhaps suggesting that thymic selection may limit the mechanistic effects of locus variation. Interestingly, we noted that the extent of N-insertion varies by ancestry-informative PCA cluster. Specifically, we found that the “Asian”-associated PCA cluster had significantly fewer N-insertions for productive TCRs when compared to the population mean which is dominated by the “Caucasian”-associated PCA cluster. This finding is, perhaps, related to the influence of broad heritable factors biasing the extent of N-insertions.

More work is required to elucidate the mechanistic relationship between *DCLRE1C* locus variation and nucleotide trimming changes. Future work can also focus on identifying correlations between TCR repertoires and host immune exposures while accounting for genetically determined repertoire biases identified here. These directions would allow us to continue disentangling the genetic and environmental determinants governing TCR repertoire diversity.

One limitation of our approach is the possibility that the SNP array data used here does not capture all potential causal variation. Further, the lack of overlap between SNP sets for the discovery and validation cohorts limited our ability to directly validate our strongest inferences. Another key constraint is the challenge of inferring the V(D)J rearrangements from the final nucleotide sequences. Therefore, there is the potential for biases resulting from incorrect V(D)J -gene assignment. We have found that controlling for D-gene assignment ambiguity in the nucleotide trimming and N-insertion analyses results in similar significant associations within the *DNTT* and *DCLRE1C* loci. Although we cannot rule out some effect of incorrect V(D)J -gene assignment bias for *trans* associations resulting from the signal being “masked” by stronger *TCRB* locus signals, these biases seem to be mostly restricted to *cis* associations.

In summary, we have found that the usage of *TCRB* genes is associated with variation in MHC and *TCRB* loci, the number of N-insertions is associated with *DNTT* variation, and the extent of nucleotide trimming is associated with *DCLRE1C* variation. Our results clearly demonstrate how variation in V(D)J recombination-related genes can bias TCR repertoire combinatorial and junctional diversity. In the case of B cells, genetically determined V(D)J gene usage biases within B-cell receptor repertoires have been linked to functional consequences for the overall immune response to specific antigens and, thus, an increased susceptibility to certain diseases (***Mikocziova et al., 2021***). As such, the genetic TCR repertoire biases identified here lay the groundwork for further exploration into the diversity of immune responses and disease susceptibilities between individuals. Such studies will enhance our understanding of how an individual’s diverse TCR repertoire can support a unique, robust immune response to disease and vaccination. Our findings also provide a step towards the ability to understand and predict an individual’s TCR repertoire composition which will be critical for the future development of personalized therapeutic interventions and rational vaccine design.

## Methods and Materials

### Discovery cohort dataset

TCR*β* repertoire sequence data for 666 healthy bone marrow donor subjects was downloaded from the Adaptive Biotechnologies website using the link provided in the original publication (***Emerson et al., 2017***). For both the discovery and validation cohorts, V, D, and J genes were assigned by comparing the TCR*β*-chain (and TCR*α*-chain for the validation cohort) nucleotide sequences to the human IMGT/GENE-DB *TCRB* (or *TCRA*) allele sequences (***Giudicelli et al., 2005***). To infer the extent of nucleotide trimming, N-insertion, and P-addition for each TCRj*β*-chain (and TCR*α*-chain) nucleotide sequence, the most parsimonious V(D)J recombination scenario was assigned to each sequence using the TCRdist pipeline (***Dash et al., 2017***). The V(D)J recombination scenario requiring the fewest N-insertions was defined as the most parsimonious scenario.

SNP array data corresponding to 398 of these subjects was downloaded from The database of Genotypes and Phenotypes (accession number: phs001918). Details of the SNP array dataset, genotype imputation, and quality control have been described previously (***Martin et al., 2020***).

### Validation cohort dataset

Peripheral blood mononuclear cell (PBMC) samples were collected from 150 healthy subjects recruited at the Health Center Sócrates Flores Vivas (HCSFV) in Managua, Nicaragua (***Ng et al., 2016***). Healthy participants were recruited as contacts of influenza infected index patients and blood samples were collected at both the initial visit and a 30 day follow-up visit. Participants provided written informed consent and parental permission was obtained from parents or legal guardians of children, in addition to verbal assent from children aged six years and older. This study was approved by the Institutional Review Boards at the University of Michigan, Centro Nacional de Diagnóstico y Referenda (Ministry of Health, Nicaragua), and University of California, Berkeley.

With these samples, PBMCs were stained with CD3-PerCP eFluor710 (Thermo Cat. 46-0037-42), CD4-BV650 (BD Biosciences Cat. 563875), CD8-APC Fire750 (Biolegend Cat. 344746), and gdCy7 (Biolegend Cat. 331222). Briefly, after thawing from cryopreservation and plating in a 96-well round bottom plate, cells were spun down and resuspended in 50 *μL* of human Fc block (BD Biosciences Cat. 564220) in Dulbecco’s phosphate-buffered saline (DPBS) at 1 *μ*L per test (1 test = 1.0 × 10^6^ cells) and incubated for 10 minutes at room temperature. Afterwards, 50 *μ*L of a Live/Dead Aqua (Tonbo Cat. 13-0870-T100, 1 *μ*L per test, 1 test = 1.0 × 10^6^ cells) and pre-titrated surface antibody cocktail in DPBS were added to each well and cells were incubated for 30 minutes on ice and in the dark. Cells were washed, resuspended in sort buffer and bulk sorted into polystyrene tubes. Afterwards, samples were spun down, pellets were resuspended in 350 *μ*L of RNA lysis buffer, and stored at −80 C in labeled epitubes.

From here, DNA was extracted from 200 *μ*L of neutrophil pellets using the Qiagen QIAamp DNA Mini Kit (Cat. 51306). Bulk repertoires for sorted CD4 and CD8 T cells were generated in accordance with the protocol developed by ***Egorov et al.*** (***2015***), and sequencing was performed on the NovaSeq by the Hartwell Center at St. Jude. Raw cDNA sequencing data were processed with the MIGEC software package (***Shugay et al., 2014***) to define error-corrected TCRA and TCRB transcript sequences, which were then analyzed as described above for the discovery cohort data (***Emerson et al., 2017***).

Genotypes for SNPs of interest corresponding to 94 of these subjects were pulled from Infinium Global Screening Array-24 v3.0 BeadChip results, which measures 654,027 SNP makers including multi-ethnic genome-wide content, curated clinical research variants, and quality control markers. High quality DNA was extracted using the Qiagen QIAamp DNA Mini Kit (Cat. 51306), and submitted to the St. Jude Hartwell Center for preparation and processing. Two SNPs, rs72640001 and rs72772435, were not included on this chip and were determined using Thermo Fisher TaqMan SNP Genotyping Assays (Cat. 4351379, Assay ID C__99271581_10 and C__99587751_10, respectively) and TaqMan Genotyping Master Mix (Cat. 4371353) according to the kit manual.

### Data preparation

With these paired SNP array and TCR-immunosequencing for both the discovery and validation cohorts, we aimed to identify significant associations between these SNPs and various TCR repertoire features. Because we would expect a difference in these phenotypes depending on whether a TCR sequence is classified as productive or non-productive, we split the data based on this TCR productivity status and computed associations separately for the two groups.

We also subset the SNP data further based on several quality control metrics. We filtered the SNP array data to use only SNPs with a minor allele frequency above 0.05 in our analyses which excluded SNPs for which all subjects had the same genotype. For the discovery cohort, this filtering procedure and previous quality control (***Martin et al., 2020***) left 6,456,824 SNPs (of the original 35 million SNPs) remaining for our analyses. Only 2 SNPs from the validation cohort overlapped with this discovery cohort SNP set. For each of these discovery and validation cohort SNPs, when fitting each association model, we excluded observations which contained a missing SNP genotype. Next, for the TCR repertoire data, we excluded repertoires which contained a relatively small number of TCRs (log_10_(TCR count) < 4.25 for productive TCRs and log_10_(TCR count) < 3.5 for non-productive TCRs) from the analyses. Lastly, for TCR*β*-chains, if a D-gene is trimmed so much that the D-gene is unidentifiable, the inference pipeline used to infer *TCRB* genes for each sequenced TCR does not report a D-gene. Instead, this D-gene (if it is indeed present) is reported as a V-J N-insertion. Because of this, we excluded these observations when fitting models for TCR features involving the D-gene (i.e. D-gene usage, both V-D and D-J junction N-insertions, D-gene P-additions, and D-gene nucleotide trimming).

### Notation

The discovery dataset contains observations for a total of *I* = 398 subjects and the validation dataset contains observations for a total of *I* = 94 subjects. Within each cohort, for subject *i* ∊ {1, …, *I*}, we observe a total of *N_i_* TCRs which, here, represents the number of TCRs which compose each subject’s TCR repertoire. Thus, for each TCR *k* ∊ {1,…, *N_i_*}, we measure a TCR feature of interest, *y_ik_*, such as the number of V-D N-insertions, the extent of V-trimming, etc. We also have SNP genotype data for a total of *J* SNPs such that for each SNP *j* ∊ {1,…, *J*} and subject *i* ∊ {1,…, *I*}, we measure the number of minor alleles in the genotype, *x_ij_* ∊ {0, 1, 2}.

### Quantifying the association strength between each SNP and TCR feature using the “simple model”

We first describe what we call the “simple model”. We will describe more complex models, as well as each model with added correction for population-substructure-related effects, in the sections following.

We began by calculating the average occurrence of the TCR feature of interest, *ȳ_i_*, within the repertoire of each subject, *i*. By condensing the data in this way, for each subject *i* ∊ {1, …, *I*}, we are left with *N_i_* = 1 observations. For example, for the discovery cohort, we can fit the model across 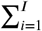 *N_i_* = 398 observations. Using this condensed dataset, for each SNP, TCR feature, and productivity status, we can fit the model:

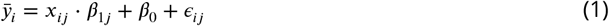

where *β*_1*j*_ is the allele effect for SNP *j* on the TCR feature of interest *ȳ_i_*, *β*_0_ is the intercept, and *ε_ij_* is the random error for subject *i* and SNP *j* such that 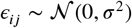.

To estimate each regression coefficient, we solved the least squares problem:

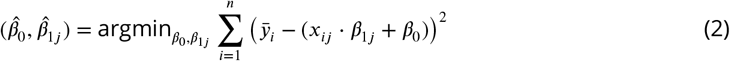

using the function `lm` in R. With each estimate of the *j*-th SNP effect on the TCR feature of interest, 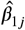, generated by fitting the least squares problem (***Equation 2***), we quantified the association strength between each SNP and the TCR feature of interest by testing whether 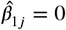. To do this, we calculate the test statistic:

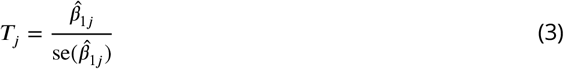

and compare *T_j_* to a *N*(0,1) distribution to obtain each P-value.

### Quantifying the association strength between each SNP and TCR feature, conditional on *TCRB* gene type using the “gene-conditioned model”

We noted that the amount of certain TCR features (such as the extent of all types of nucleotide trimming) vary by V(D)J *TCRB* gene choice. Thus, we can condition on this gene choice to quantify the direct association between each SNP and the amount of each TCR feature, without confounding gene choice effects. In this way, we condition on each gene type *t* ∊ {V-gene,J-gene, D-gene}corresponding to the TCR feature of interest (i.e. *t* = V-gene for V-gene trimming, *t* = J-gene for J-gene trimming, etc.). We will refer to the following model as the “gene-conditioned model” in the main text. Many similarities exist between the “simple model” described in the previous section and this “gene-conditioned model.” Thus, we will focus on the differences between the two models here. We will describe both models with added correction for population-substructure-related effects, in the sections following.

As in the previous section, we, again, want to reduce the number of data observations. For each subject *i* ∊ {1, …, 1}, we can calculate the average amount of each TCR feature *ȳ_im_* by each candidate *TCRB* gene allele group *m* for the given gene type *t* such that *m* ∊ {1, …, *M_t_*}. In calculating the average amount of each TCR feature across TCRs with the same candidate *TCRB* gene allele, we combined *TCRB* gene alleles which had identical CDR3 sequences and were of the same candidate *TCRB* gene into *TCRB* gene allele groups. As such, the number of observations per subject *N_i_* in this condensed dataset will equal *M_t_* and, thus, we will need to fit each model across 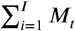 observations. In our data, for TCR*β* chains, we observe 141 possible *TCRB* V-gene allele groups, 16 J-gene allele groups, and 3 D-gene allele groups. Thus, using the extent of nucleotide trimming as an example TCR feature within the discovery cohort, with this condensed formulation, for each SNP and productivity status, we have ~ 56,000 observations for V-gene trimming, ~ 6,000 observations for J-gene trimming, and ~ 1,200 observations for both types of D-gene trimming.

Using this condensed dataset, for each SNP, TCR feature, and productivity status, we fit the following “gene-conditioned model”:

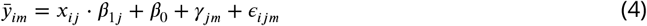

where *γ_jm_* represents the gene-effect on the amount of the TCR feature of interest for SNP *j* and gene-allele-group *m*, and *ε_ijm_* is the random error for subject *i*, SNP *j*, and gene-allele-group *m* such that 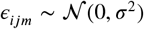. The variables *x_ij_*, *β*_1*j*_, and *β*_0_ are defined as in the “simple model” description (***Equation 1***) in the previous section. However, since each subject had a different number of TCRs measured and varying *TCRB* gene usage, we calculated the proportion of TCRs from each candidate *TCRB* gene allele group, *m*, to define a weight, *W_im_*, for each observation:

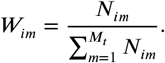

With this, we solved the following weighted least squares problem for each SNP, TCR feature, and productivity status combination:

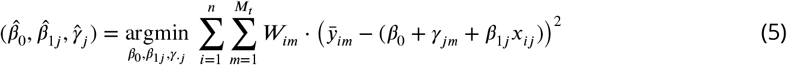

using the `lm` function in R.

With each estimate of the *j*-th SNP effect on the amount of the TCR feature of interest, 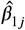, generated using the models described above, we quantified the association strength between each SNP and the amount of the TCR feature by testing whether 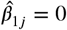. To do this, we applied a t-test (described in the previous section) using the test statistic (***Equation 3***) to obtain each P-value. However, because our condensed dataset contains a total of *M_t_*, observations from each subject *i*, these P-values may be inflated due to intra-subject observations being potentially correlated. Thus, to increase the accuracy of the P-value calculation, for each association P-value below a certain threshold (we chose *P* < 5 × 10^−5^), we recalculated the P-value using a clustered bootstrap (with subjects as the sampling unit). To do so, for each bootstrap iterate, we resampled subjects from the condensed dataset with replacement. Using this re-sampled data, we fit the model in ***Equation 5*** to estimate each coefficient. We repeated this bootstrap process 100 times and used the resulting 100 coefficient estimates to estimate a standard error for each model coefficient. With this re-calculated standard error of the estimate of the *j*-th SNP effect on the amount of the TCR feature of interest, 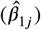, we wanted to test whether 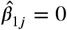 by recalculating the test-statistic, ***Equation 3***, and applying a t-test to obtain each “corrected” P-value. As noted in the multiple testing correction methods section, when accounting for multiple testing via Bonferroni correction, we used the entire number of TCR features and SNPs considered (not just those that were sufficiently promising to warrant use of the bootstrap to get a more accurate P-value): This ensures that our correction will not be anti-conservative.

### Correcting for population-substructure-related effects

Structure within our SNP genotype data (such as population-substructure-related biases due to ancestry), if present, may produce false positive associations when quantifying the association strength between each SNP and our phenotype of interest. To account for this, we implemented principal component analysis as commonly applied to genome-wide genotype data for population substructure inference. Specifically, we used the PC-AiR algorithm (***Conomos et al., 2015***) which identifies principal components that capture ancestry while accounting for relatedness in the samples. As such, the top principal components calculated from the genotype data reflect population substructure among the samples. When plotting the proportion of variance explained by each PC, we find that the majority of variability appears to be explained by the top eight PCs (***Figure 9***). This conclusion is supported when plotting each PC score by ancestral group (***Figure 9***). With this, we incorporated the top eight principal components as covariates into our GWAS models described above.

**Figure 9.**
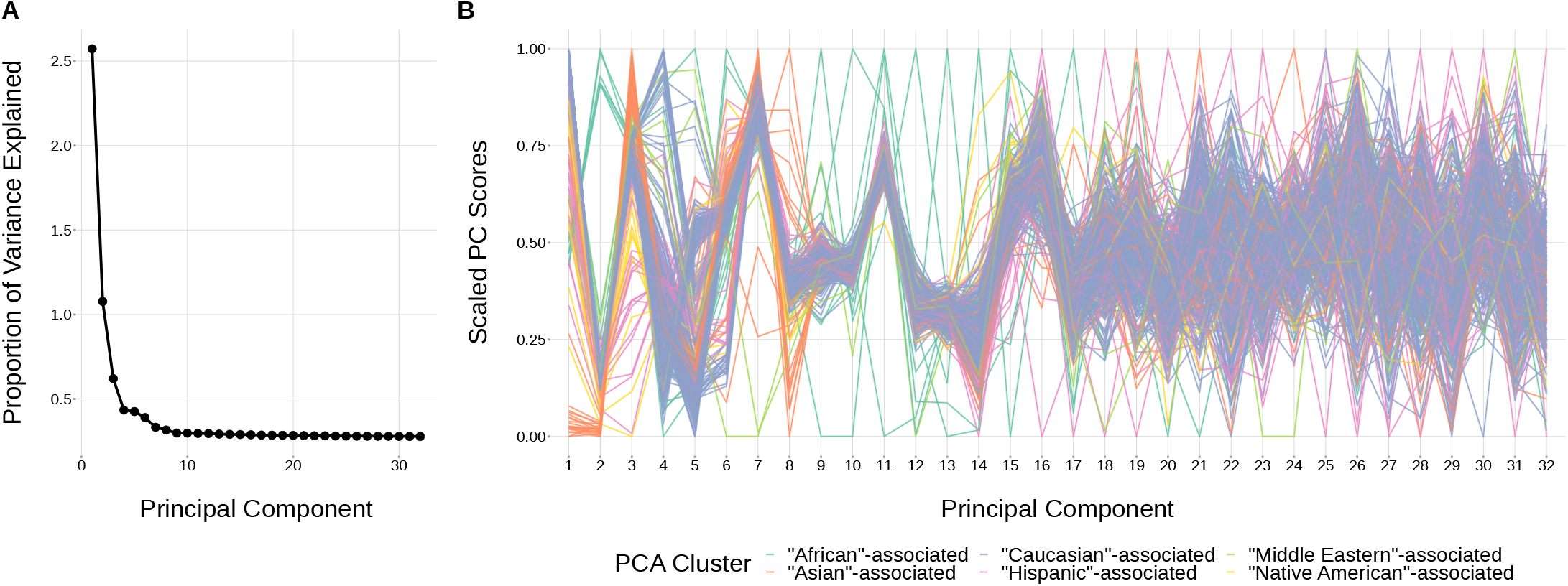
The top principal components calculated from genotype data reflect ancestry structure among samples. (**A**) The majority of the ancestry-informative principal component analysis variance is explained by the first 8 principal components. (**B**) The first 8 principal components show distinct separation by PCA cluster. Each colored line represents one of the 398 samples. The first 32 principal components are shown on the X-axis and their scaled component values for each subject on the Y-axis.

As such, to quantify the association strength between each SNP and TCR feature without conditioning on gene usage as in ***Equation 1***, while incorporating principal component terms to correct for population-substructure-related bias due to ancestry, we fit the model:

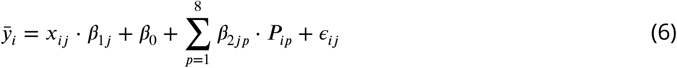

where *ȳ_i_*, *x_ij_*, *β*_1*j*_, *β*_0_, and *ε_ij_* are defined as in ***Equation 1***, *β*_2*jp*_ is the population-substructure-related bias correction term for SNP *j* and the *p*-th principal component, and *P_ip_* is the *p*-th principal component for subject *i* as calculated above. To estimate each regression coefficient, we solved the following least squares problem for each SNP, TCR feature, and productivity status combination:

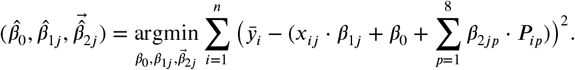

Furthermore, to quantify the association strength between each SNP and TCR feature, conditional on gene usage as in ***Equation 4***, while incorporating principal component terms to correct for population-substructure-related bias due to ancestry, we fit the model:

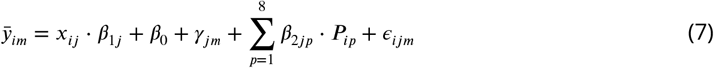

where *ȳ_im_*, *x_ij_*, *β*_1*j*_, *β*_0_, *γ_jm_*, and *ε_ij_* are defined as in ***Equation 4*** and *β*_2*jp*_ and *P_ip_* are defined as in ***Equation 6***. Again, to estimate each regression coefficient, we solved the following weighted least squares problem for each SNP, TCR feature, and productivity status combination:

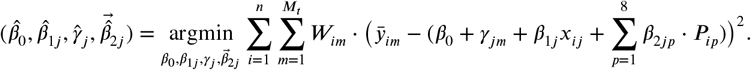

With these estimates for the population-substructure-corrected *j*-th SNP effect on the amount of the TCR feature of interest, 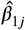, we calculated a P-value using the methods described in the methods section for each model type.

### Correcting for *TRBD2* allele genotype, SNP genotype linkage when quantifying SNP, TCR feature associations within the *TCRB* locus

Within the *TCRB* locus, we noted that SNP genotypes were associated with *TRBD2* allele genotype (***Figure 3-Figure Supplement 1***). Associations between gene-alleles and *TCRB* locus SNP genotypes, if present, may produce false positive associations when implementing the “gene-conditioned model” to infer associations between SNPs and TCR repertoire features, conditional on gene usage. To explore this phenomenon further, we zoomed in to the *TCRB* locus and incorporated a *TRBD2* allele genotype correction procedure into our model formulation. As such, to quantify the association strength between each *TCRB* locus SNP and TCR feature, conditional on gene usage and correcting for population-substructure-related effects as in ***Equation 7***, while incorporating *TRBD2* allele genotype correction terms, we fit the model:

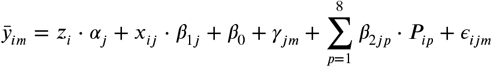

where *z_i_* represents the qualitative *TRBD2* allele genotype status for subject *i* such that *z_i_* ∊ {“TRBD2*01 homozygous”, “heterozygous”, “TRBD2*02 homozygous”}, *α_j_* is the *TRBD2* allele genotype effect for SNP *j*, and the remaining variables are defined as in ***Equation 7***. With this model formulation, we can estimate each regression coefficient by solving the following weighted least squares problem for each *TCRB* SNP, TCR feature, and productivity status combination:

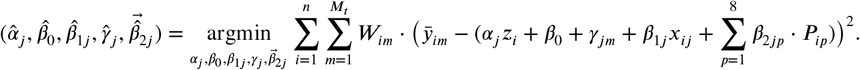

With these estimates for the *TRBD2* allele genotype and population-substructure-corrected *j*-th SNP effect on the amount of the TCR feature of interest, 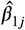, we calculated a P-value using the methods described in the methods section for the “gene-conditioned model”.

### Multiple testing correction for associations

For each TCR feature (i.e. extent of trimming, number of N-insertions, etc.), we considered the significance of associations using a Bonferroni-corrected threshold. To establish each threshold, we corrected for each TCR feature subtype (i.e. V-gene trimming, J-gene trimming, etc. for the TCR trimming feature), the two TCR productivity types (productive and non-productive), and the total number of SNPs tested. When considering associations in the whole-genome context, we corrected for the approximately 6.5 million SNPs (remaining after filtering). When considering associations in a gene-level context, we corrected for the number of SNPs within 200 kb of the gene of interest. For the validation analysis, we considered associations in a SNP-level context and did not correct for multiple SNPs. However, for the validation analysis, we considered the significance of associations within both TCR*α* and TCR*β* chains and, thus, corrected the significance threshold accordingly.

### Genomic inflation factor calculations

We defined the genomic inflation factor *λ* to be the ratio of the median of the empirically observed squared test statistic to the expected median (***Devlin and Roeder, 1999**; **Freedman et al., 2004**; **Price et al., 2010***). For each GWAS analysis implemented using the “simple model”, we used the test statistic *T_j_* given by ***Equation 3*** for each SNP *j* = {1 … *J*} tested genome-wide. For each GWAS analysis implemented using the “gene-conditioned model”, it was not computational feasible to calculate a test statistic *T_j_* for all SNPs tested genome-wide using the bootstrapping protocol described in the “gene-conditioned model” methods section. Thus, instead, we randomly sampled 10,000 SNPs and calculated the test statistic *T_j_* for each SNP in the random subset. Let 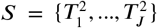 be the set of all squared test statistics. As such,

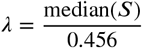

where 0.456 is the median of a chi-squared distribution with one degree of freedom. If the GWAS analysis results follow the chi-squared distribution, the expected value of *λ* is 1. Thus, when *λ* < 1.03, we concluded that there was no evidence of systemic population-substructure-related bias in the analysis (***Price et al., 2010**; **Conomos et al., 2016***).

### Conditional analysis to test for multiple independent association signals

Within the *DNTT* and *DCLRE1C* loci, we performed a stepwise series of nested regression analyses to test for independent SNP associations within each locus for N-insertion and nucleotide trimming, respectively. We used the same models and covariates as the primary analyses (“simple model” for associations between N-insertion and *DNTT* variation and the “gene-conditioned model” for associations between nucleotide trimming and *DCLRE1C* variation) and included the most significant SNP within each locus as an additional covariate. We inferred the association between each SNP within each locus and the TCR feature of interest using this new conditional model and considered significant associations at a gene-level Bonferroni-corrected significance threshold for each locus. From here, we repeated this analysis (if necessary), identifying and adding additional SNPs one-by-one as a covariate to each successive model. Once the P-value of top SNP within the locus was no longer significant, we concluded the analysis. SNPs which were added as as additional covariates in the final conditional model were considered to be independent signals.

### Ancestry-informative PCA cluster classification

In order to correct for population-substructure-related biases due to ancestry in our GWAS analyses, we used ancestry-informative principal component analysis. The original genotyping dataset (***Martin et al., 2020***) contained self-reported ancestry. However, a number of subjects did not self-report ancestry in the original data collection. Further, for some subjects, their self-reported ancestry was discordant with clusters observed in a principal component analysis. Therefore, for analysis purposes, we used the minimum covariance determinant method (***Rousseeuw and Van Driessen, 1999**; **Conomos et al., 2016***) with the original self-identified labels to group the subjects into six ancestry-informative PCA clusters: “African”-associated (8), “Asian”-associated (23), “Caucasian”-associated (322), “Hispanic”-associated (30), “Middle Eastern”-associated (5), and “Native American”-associated (10).

### Quantifying associations between *TRBD2* allele genotype and SNP genotype within the *TCRB* locus

For each significantly associated SNP within the *TCRB* locus as shown in ***Figure 3***, we compared SNP genotype to *TRBD2* allele genotype across all subjects. We used Pearson correlation to measure the correlation between the two genotypes.

### Quantifying TCR repertoire feature and SNP minor allele frequency variations by ancestry-informative PCA cluster

To quantify PCA cluster variation of TCR repertoire features (such as total N-insertions (V-D N-insertion and D-J N-insertion)), we first calculated an average of each TCR repertoire feature by subject and productivity status. We also calculated a population mean of each TCR repertoire feature by productivity status. Each subject was classified into one of six PCA clusters. Thus, we compared the mean of the TCR repertoire features within each PCA cluster to the population mean using a one-sample t-test to compute each P-value. We used Bonferroni multiple testing correction to adjust each P-value.

We also calculated SNP minor allele frequencies for the whole population and for each PCA cluster individually such that

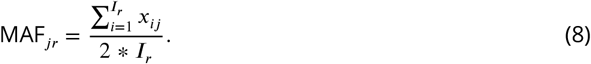

Here, MAF_*jr*_ is the minor allele frequency for SNP marker *j* and PCA cluster *r*, *I_r_* is the number of individuals in the PCA cluster *r*, and *x_ij_* is the number of alleles in the genotype of SNP marker *j* for subject *i* ∊ {1,…, *I_r_*}. For each SNP *j*, the minor allele was defined as the allele with the lowest frequency in the total population. To quantify minor allele frequency differences by PCA cluster for select SNPs within various loci of interest (i.e. *DNTT* gene), we compared the minor allele frequencies calculated within PCA-clusters to the minor allele frequencies calculated for the entire population using a one-sample t-test to compute each P-value. Again, we used Bonferroni multiple testing correction to adjust each P-value.

For both of these analyses, we used the `t_test` function from the `rstatix` package in R.

### Implementation and code

R code implementing the genome-wide association inferences described here is available at https://github.com/phbradley/tcr-gwas. The following tools were especially helpful:

- data.table (***Dowle and Srinivasan, 2021***)
- tidyverse (***Wickham et al., 2019***)
- doParallel (***Corporation and Weston, 2020***)
- SNPRelate (***Zheng et al., 2012***)
- GWASTools (***Gogarten et al., 2012***)
- GENESIS (***Gogarten et al., 2019***)
- cowplot (***Wilke, 2020***)

## Acknowledgments

The authors thank Christopher Carlson, William DeWitt, and Michael Lieber for helpful discussions regarding this paper.

## Additional Information

### Funding

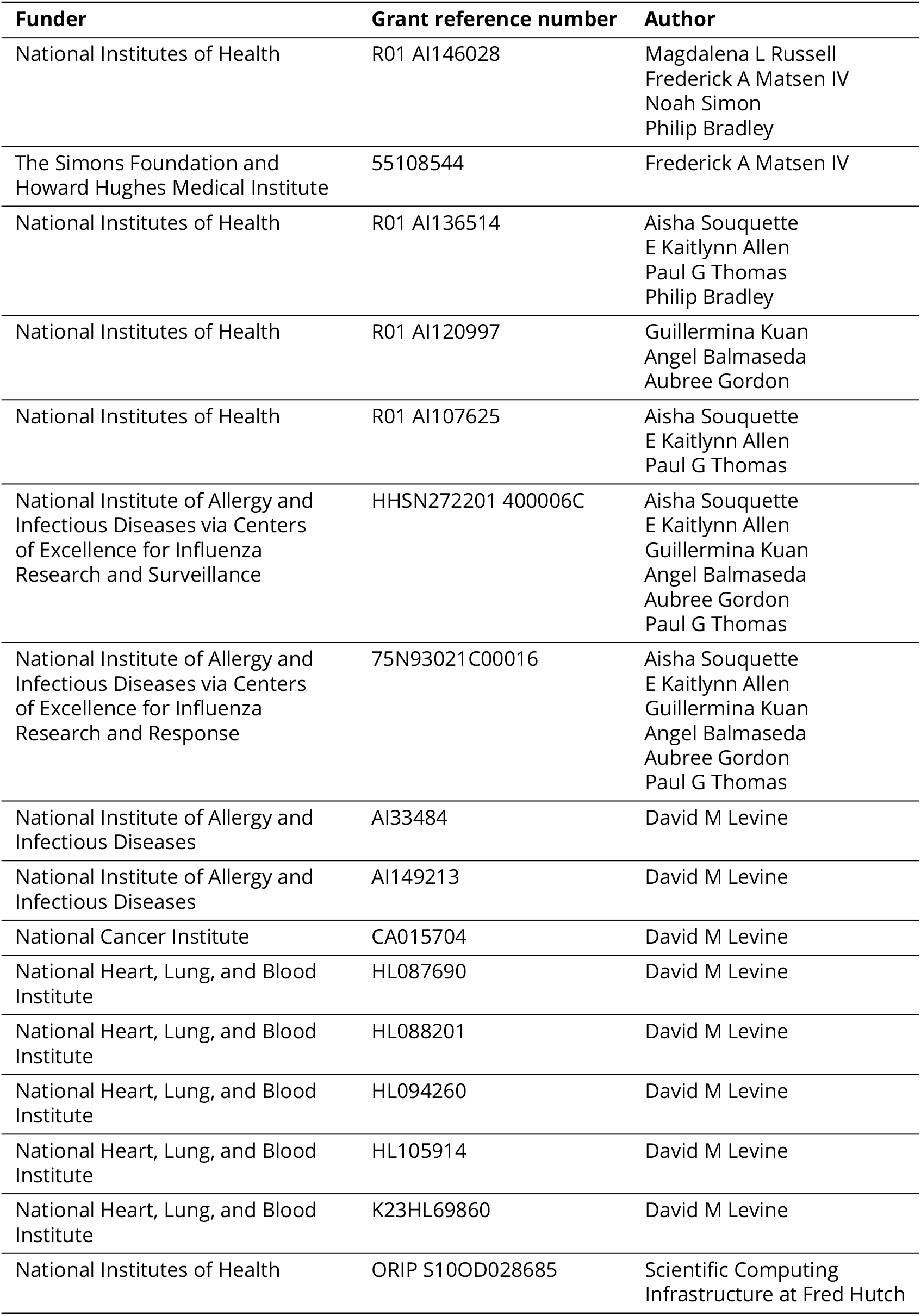

## Additional Files

### Data availability

Data required to reproduce the findings reported here and a table mapping subject identifiers between the TCR repertoire and SNP data for the discovery cohort will be deposited in the Zenodo database prior to publication.

The following new dataset was generated:

| Author | Year | Dataset title | Dataset URL | Database, license, and accessibility information |
| --- | --- | --- | --- | --- |
| Aisha Souquette, E Kaitlynn Allen, Guillermina Kuan, Angel Balmaseda, Aubree Gordon, Paul G Thomas | 2021 | The Nicaraguan Influenza Cohort Study | https://www.ncbi.nlm.nih.gov/bioproject/PRJNA762269 | Publicly available in The BioProject database (accession number: PRJNA762269) |

The following previously published datasets were used:

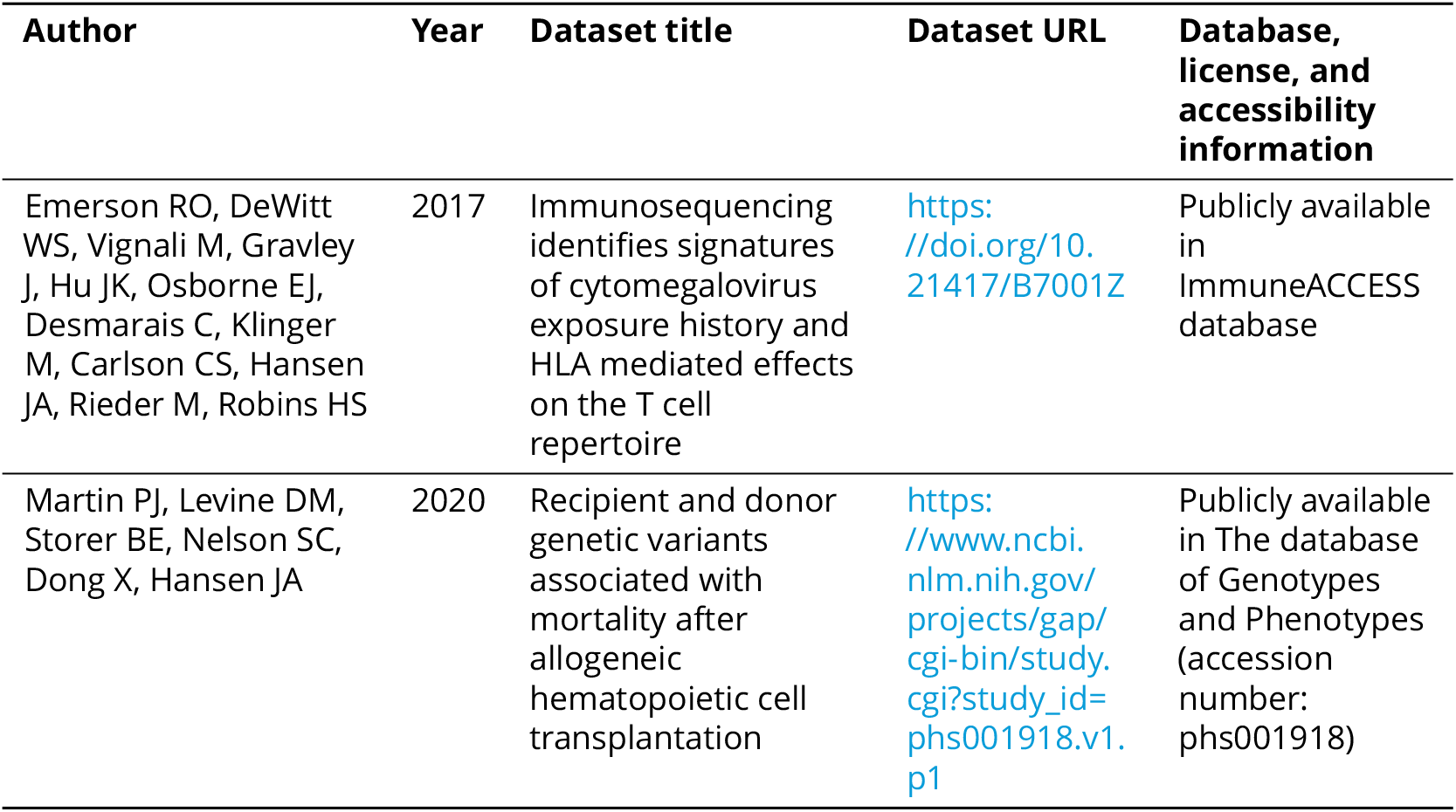

---

**Table 2-Figure supplement 1.**
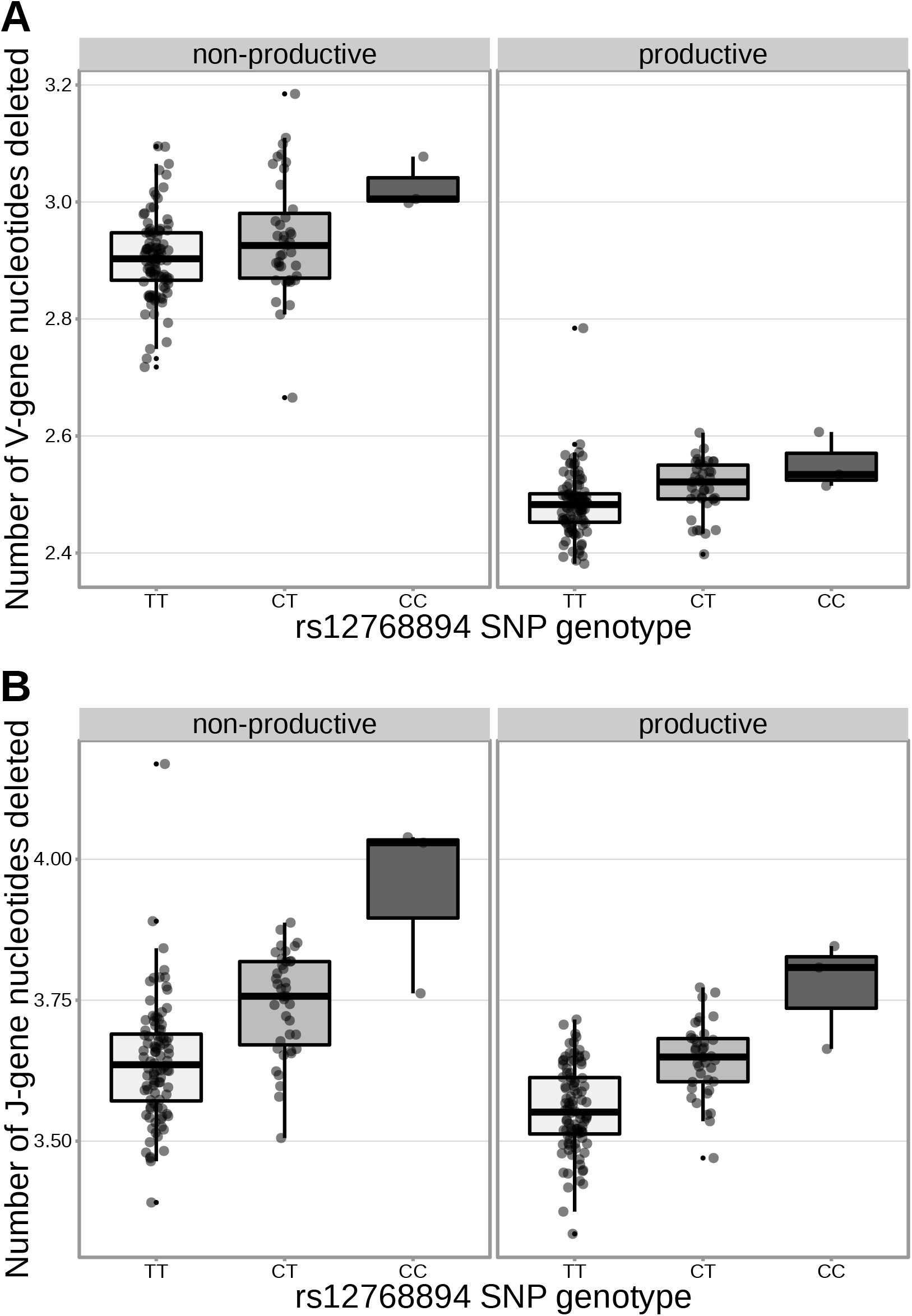
The extent of V- and J-gene trimming of productive and nonproductive *TCRβ* chains changes as a function of SNP genotype within the discovery cohort for a non-synonymous *DCLRE1C* SNP (rs12768894, c.728A>G). Only TCRs containing *TRBJ1-1*01* (the most frequently used *TRBJ1* gene across subjects) were included when calculating the average number of J-gene nucleotides deleted for each subject. Only TCRs containing *TRBV5-1*01* (the most frequently used *TRBV* gene across subjects) were included when calculating the average number of V-gene nucleotides deleted for each subject.

**Table 2-Figure supplement 2.**
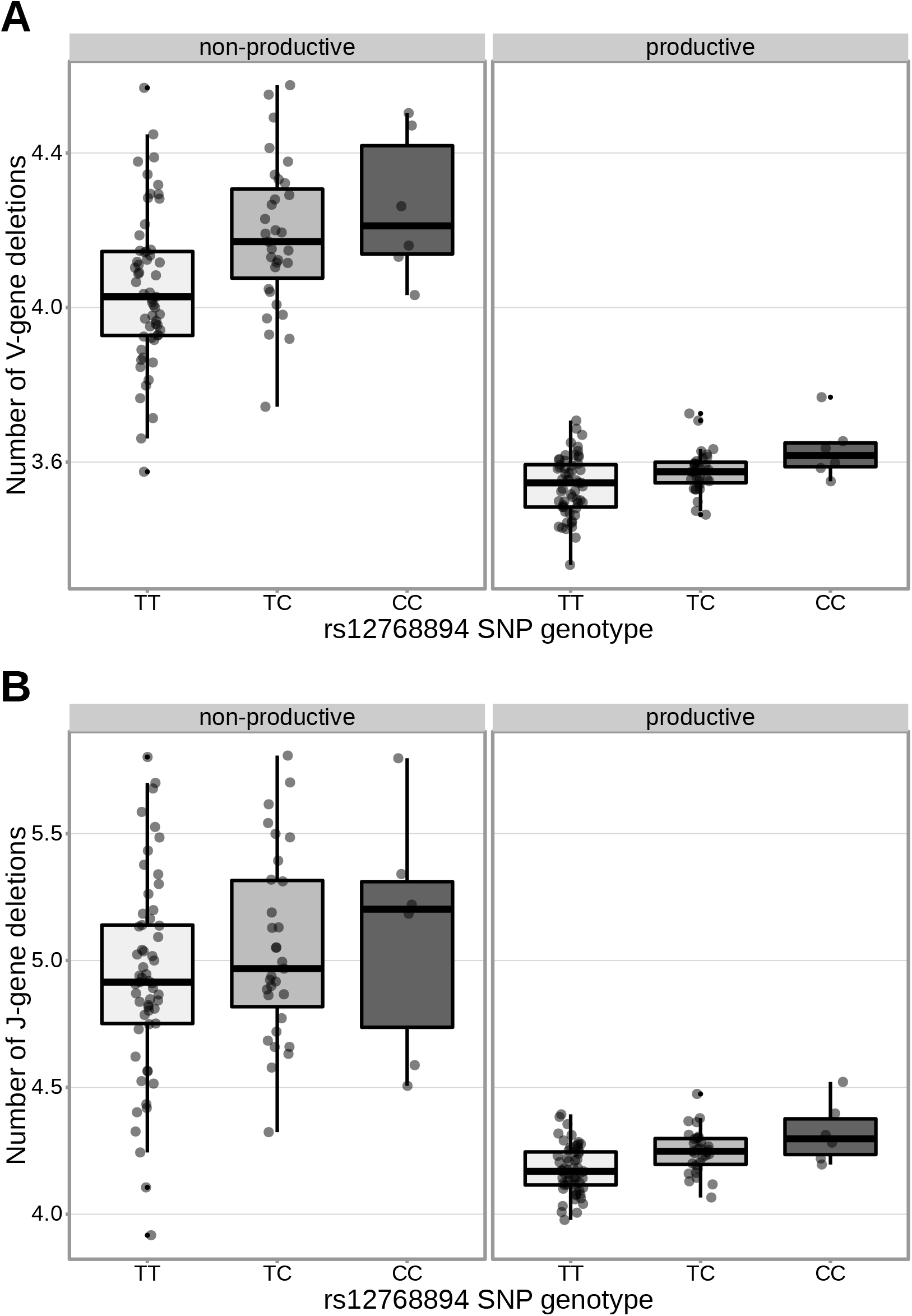
The extent of V-gene trimming (**A**) of productive and non-productive TCR*β* chains and J-gene trimming (**B**) of productive TCR*β* chains changes as a function of SNP genotype within the validation cohort for a non-synonymous *DCLRE1C* SNP (rs12768894, c.728A>G). The average number of nucleotides deleted was calculated across all TCR*β* chains for each subject, regardless of gene-usage.

**Table 2-Figure supplement 3.**
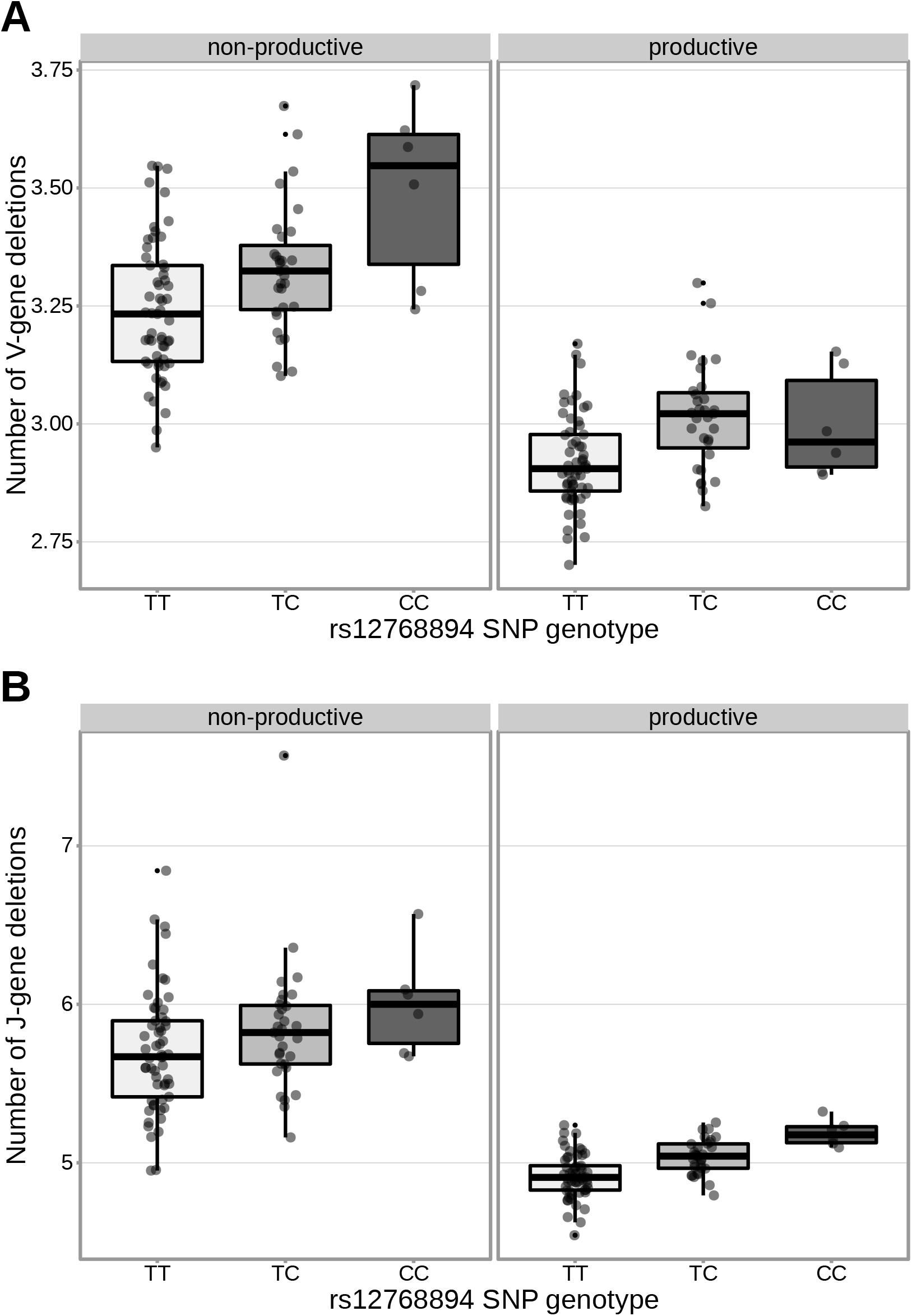
The extent of V- (**A**) and J-gene (**B**) trimming of productive and non-productive TCR*α* chains changes as a function of SNP genotype within the validation cohort for a non-synonymous *DCLRE1C* SNP (rs12768894, c.728A>G). The average number of nucleotides deleted was calculated across all TCR*α* chains for each subject, regardless of gene-usage.

**Figure 3-Figure supplement 1.**
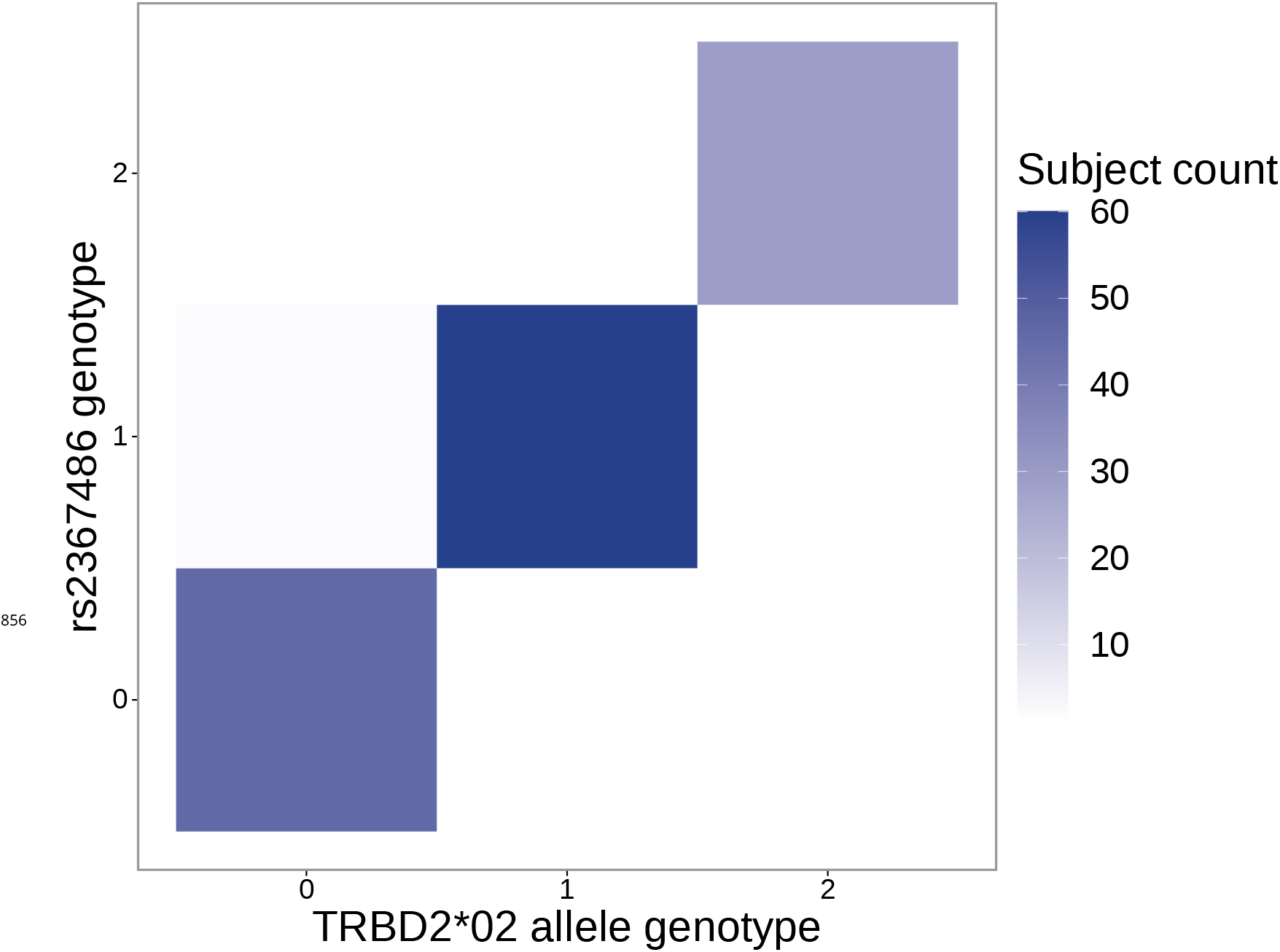
The SNP genotype for the SNP (rs2367486) most significantly associated with 5’ end D-gene trimming within the *TCRB* locus is also associated with *TRBD2*02* allele genotype. Specifically, SNP genotype and *TRBD2*02* allele genotype are significantly correlated (*P* < 2.2 × 10^−16^ and *χ*^2^ = 259.3) using a chi-square test of independence. The Y-axis integer genotypes correspond to the number of minor alleles within the rs2367486 SNP genotype. The X-axis integer genotypes correspond to the number of *TRBD2*02* alleles within the *TRBD2* gene locus genotype.

**Figure 3-Figure supplement 2.**
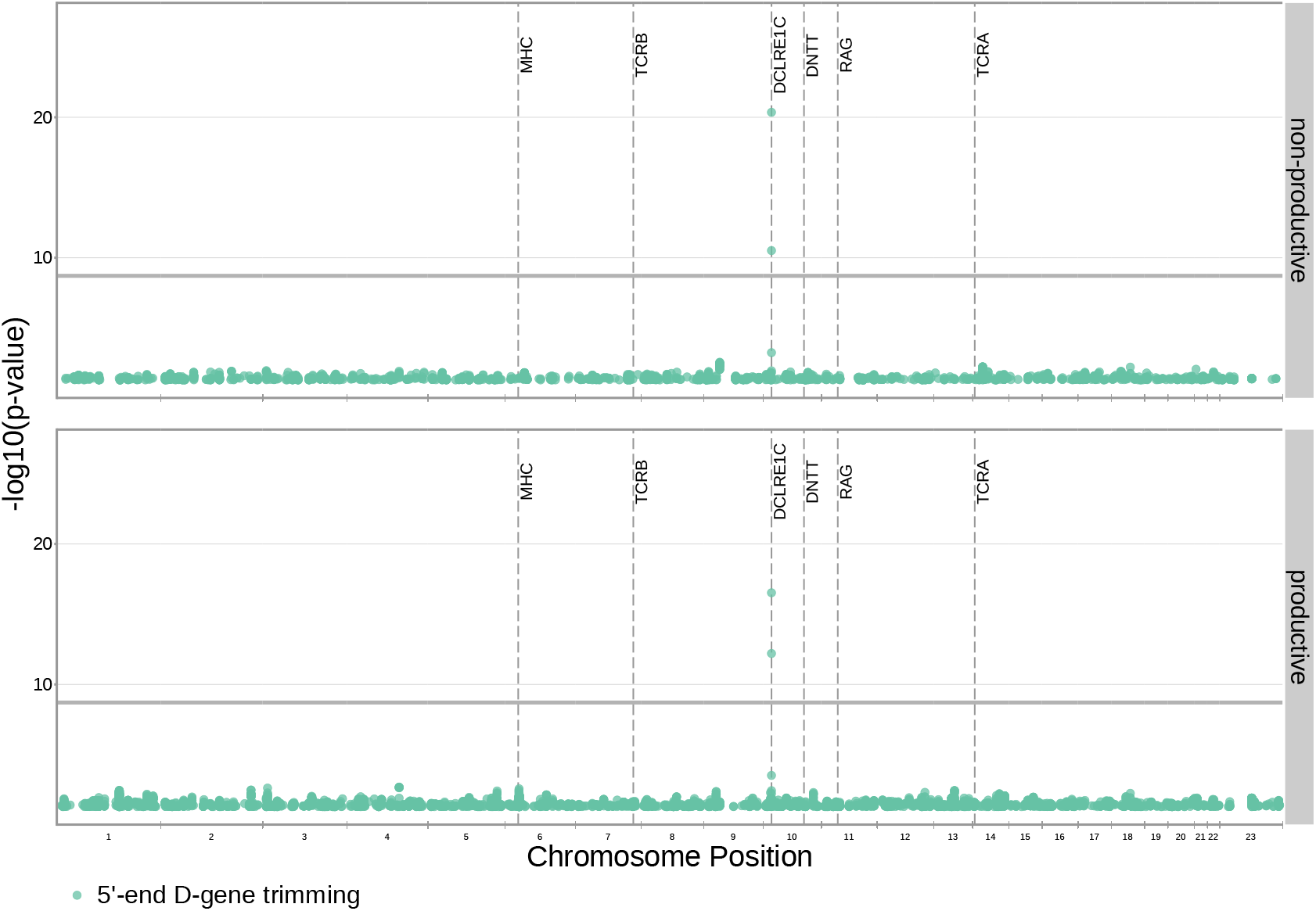
Significant associations are no longer observed between 5’ end D-gene trimming and variation in the *TCRB* locus after correcting for *TRBD2* allele genotype in our model formulation. Further, four new significant associations are present between 5′ end D-gene trimming and variation in the *DCLRE1C* locus. Only SNP associations whose *P* < 5 × 10^−2^ are shown here. All genome-wide 3′ end D-gene trimming associations fell above this plotting threshold. The gray horizontal line corresponds to a P-value of 1.94 × 10^−9^ (calculated using whole-genome Bonferroni correction, see Methods).

**Figure 3-Figure supplement 3.**
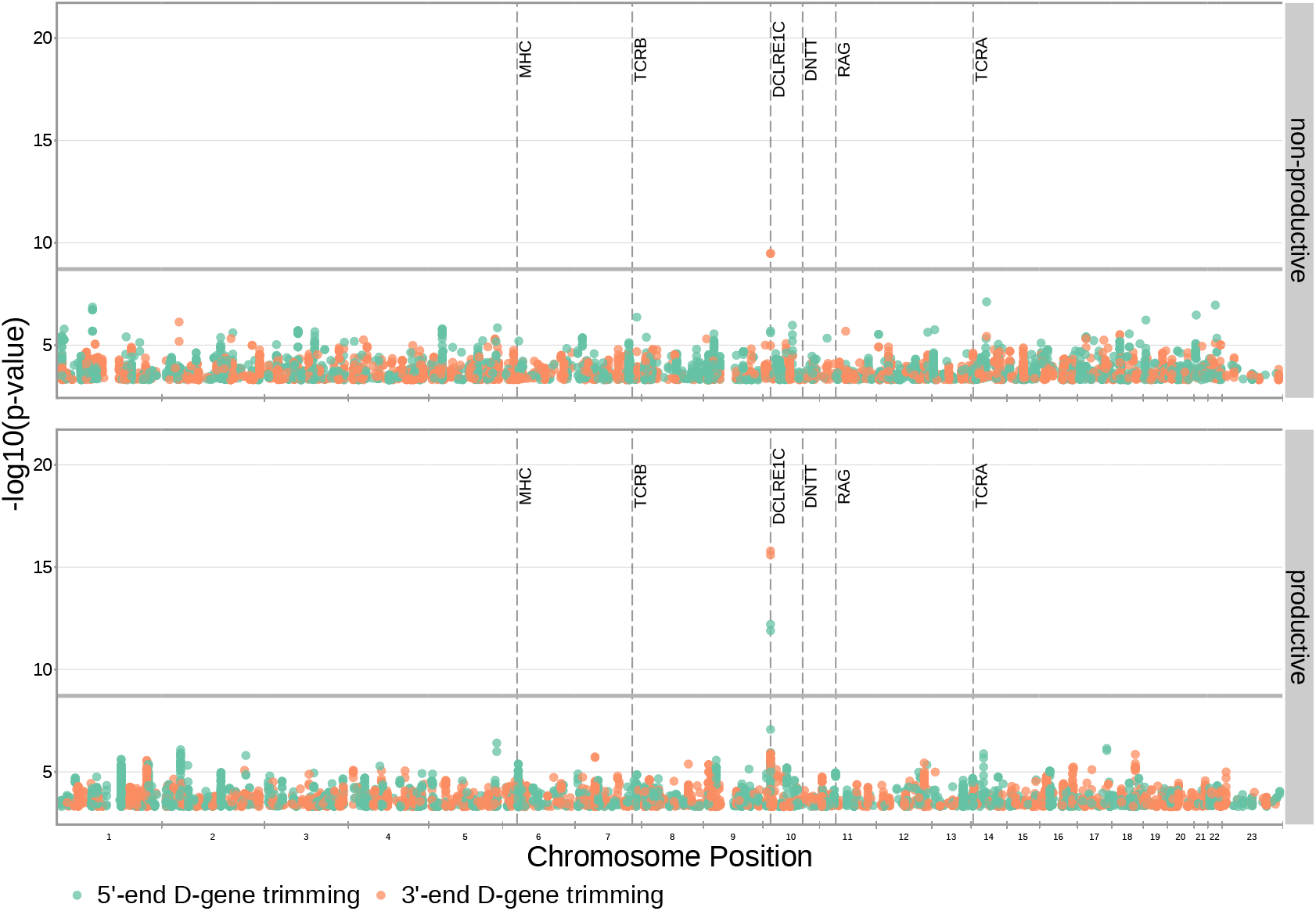
Significant associations are also no longer observed between 5’ end D-gene trimming and variation in the *TCRB* locus when restricting the analysis to TCRs which contain *TRBJ1* genes (and consequently contain *TRBD1*). Additionally, two new associations are present between 5′ end D-gene trimming and variation in the *DCLRE1C* locus for productive TCRs. Four new associations are present between 3′ end D-gene trimming and variation in the *DCLRE1C* locus. Only SNP associations whose *P* < 5 × 10^−4^ are shown here. The gray horizontal line corresponds to a P-value of 1.94 × 10^−9^ (calculated using whole-genome Bonferroni correction, see Methods).

**Figure 3-Figure supplement 4.**
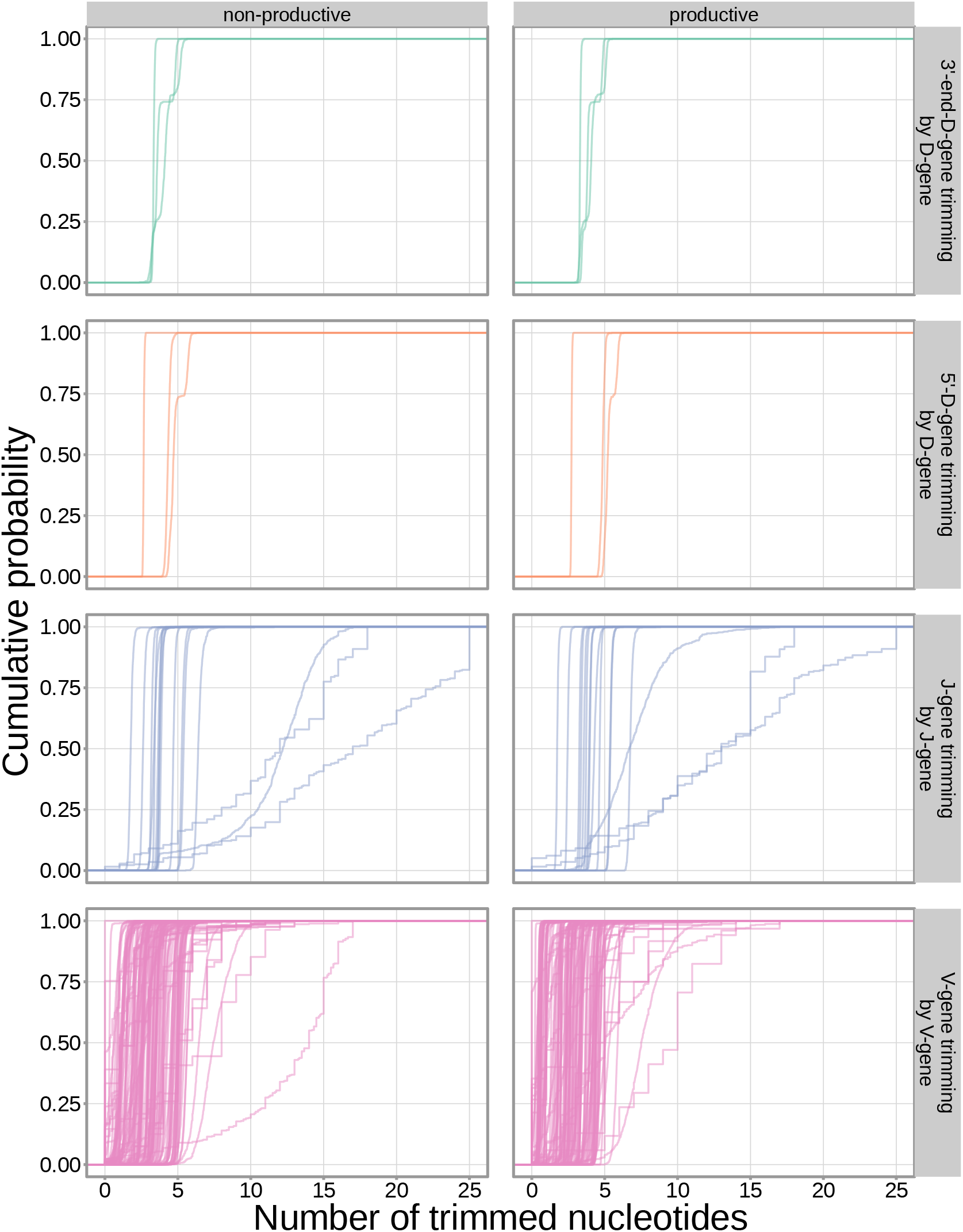
The extent of nucleotide deletion varies by the gene allele identity for all gene types. An empirical cumulative distribution is drawn for each gene allele type within each indicated gene type (i.e. V-gene, D-gene, J-gene).

**Figure 3-Figure supplement 5.**
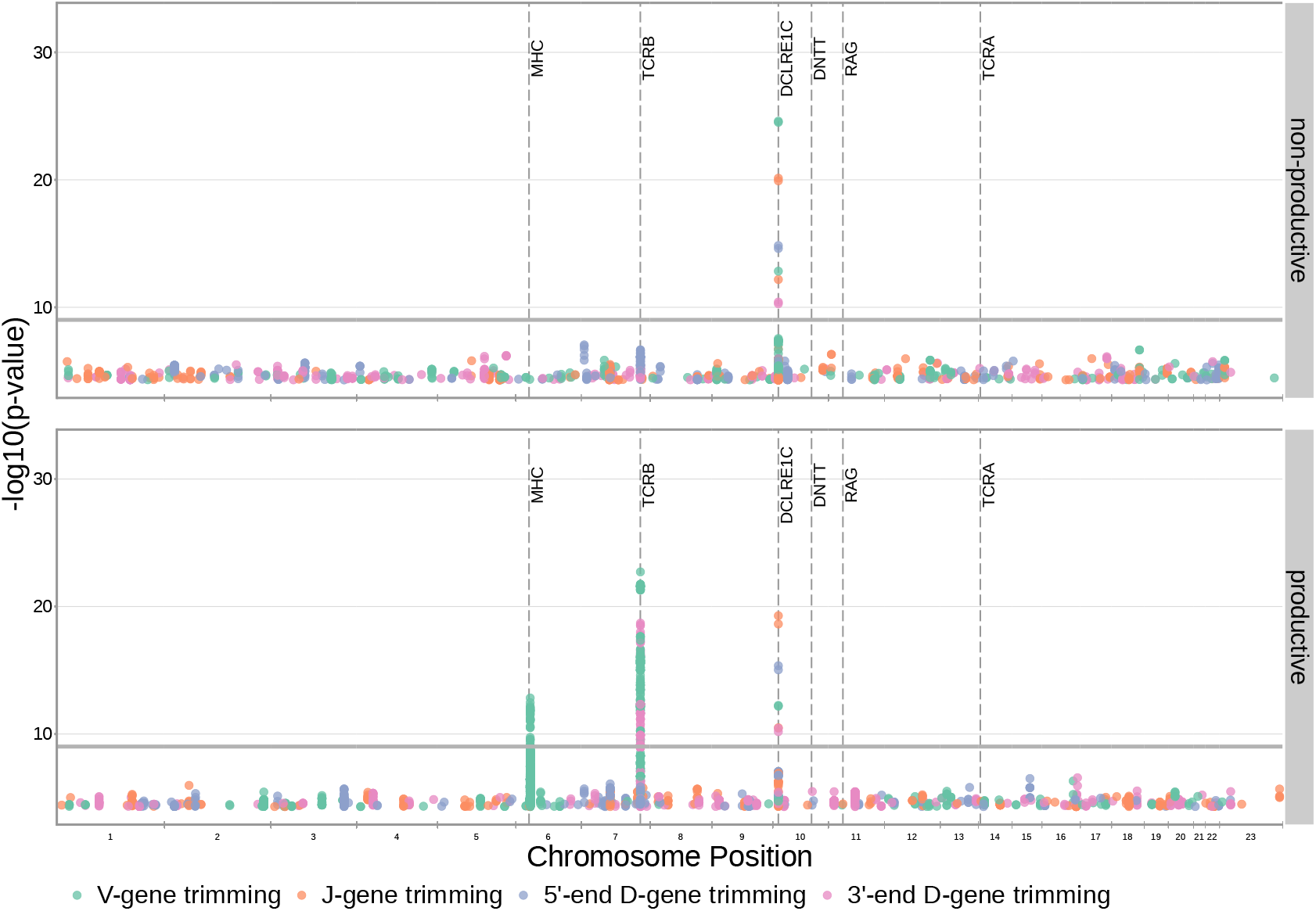
Significant SNP associations are located within the MHC, *TCRB* and *DCLRE1C* loci for all four trimming types when calculating the strength of association without conditioning out effects mediated by gene choice. Earlier findings relating variations in MHC and *TCRB* to gene usage changes, however, indicate that many of these associations are likely artefactual. Only SNP associations whose *P* < 5 × 10^−5^ are shown here. The gray horizontal line corresponds to a P-value of 9.68 × 10^−10^.

**Figure 3-Figure supplement 6.**
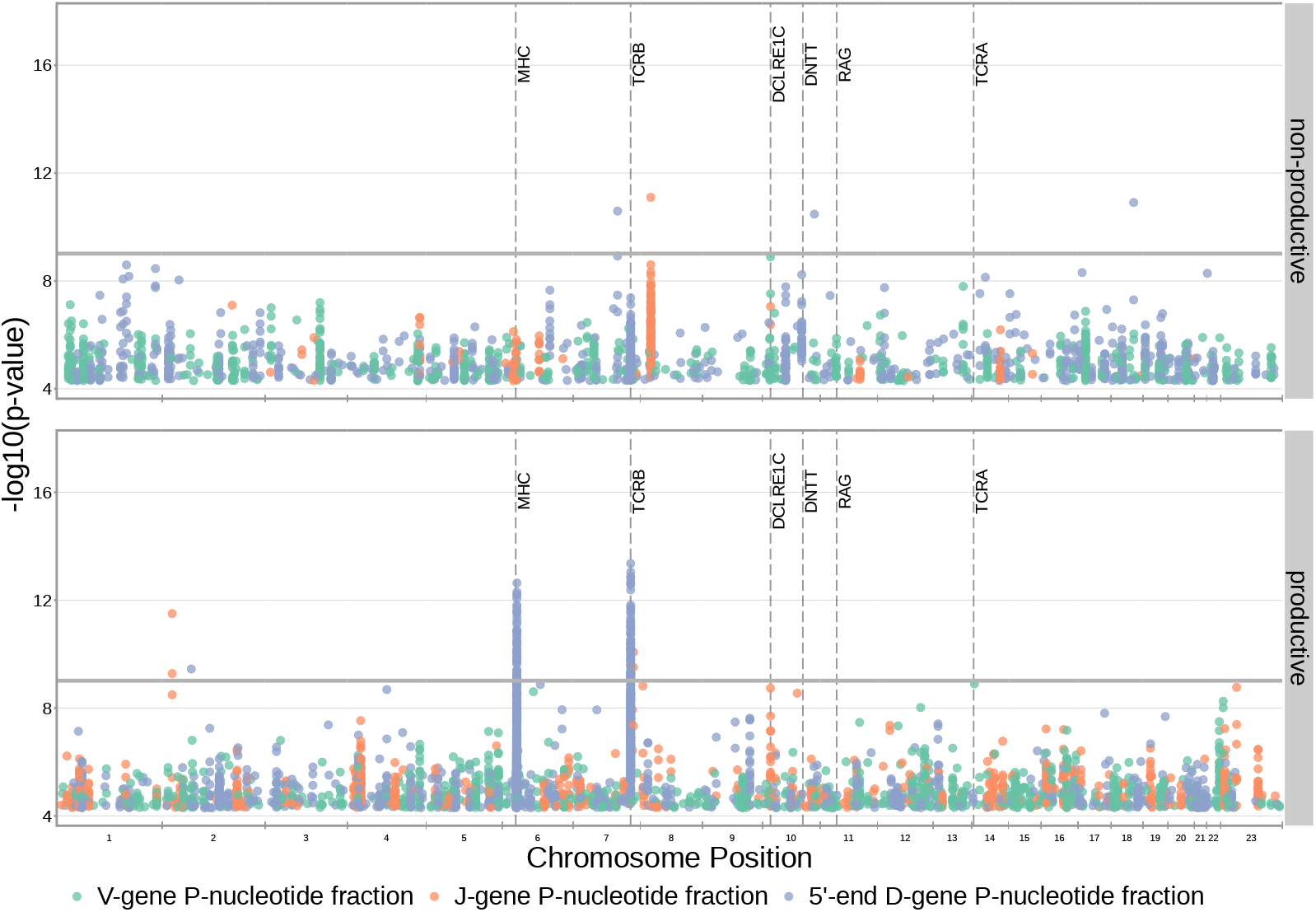
SNP associations for all fractions of non-gene-trimmed TCRs containing P-nucleotides are not significant within the *DCLRE1C* locus. However, significant associations are present within the *TCRB* and MHC loci for the fraction of non-D-gene-trimmed, productive TCRs containing 5′ end D-gene P-nucleotides. Only SNP associations whose *P* < 5 × 10^−5^ are shown here. The gray horizontal line corresponds to a P-value of 9.68 × 10^−10^ (calculated using whole-genome Bonferroni correction, see Methods).

**Figure 3-Figure supplement 7.**
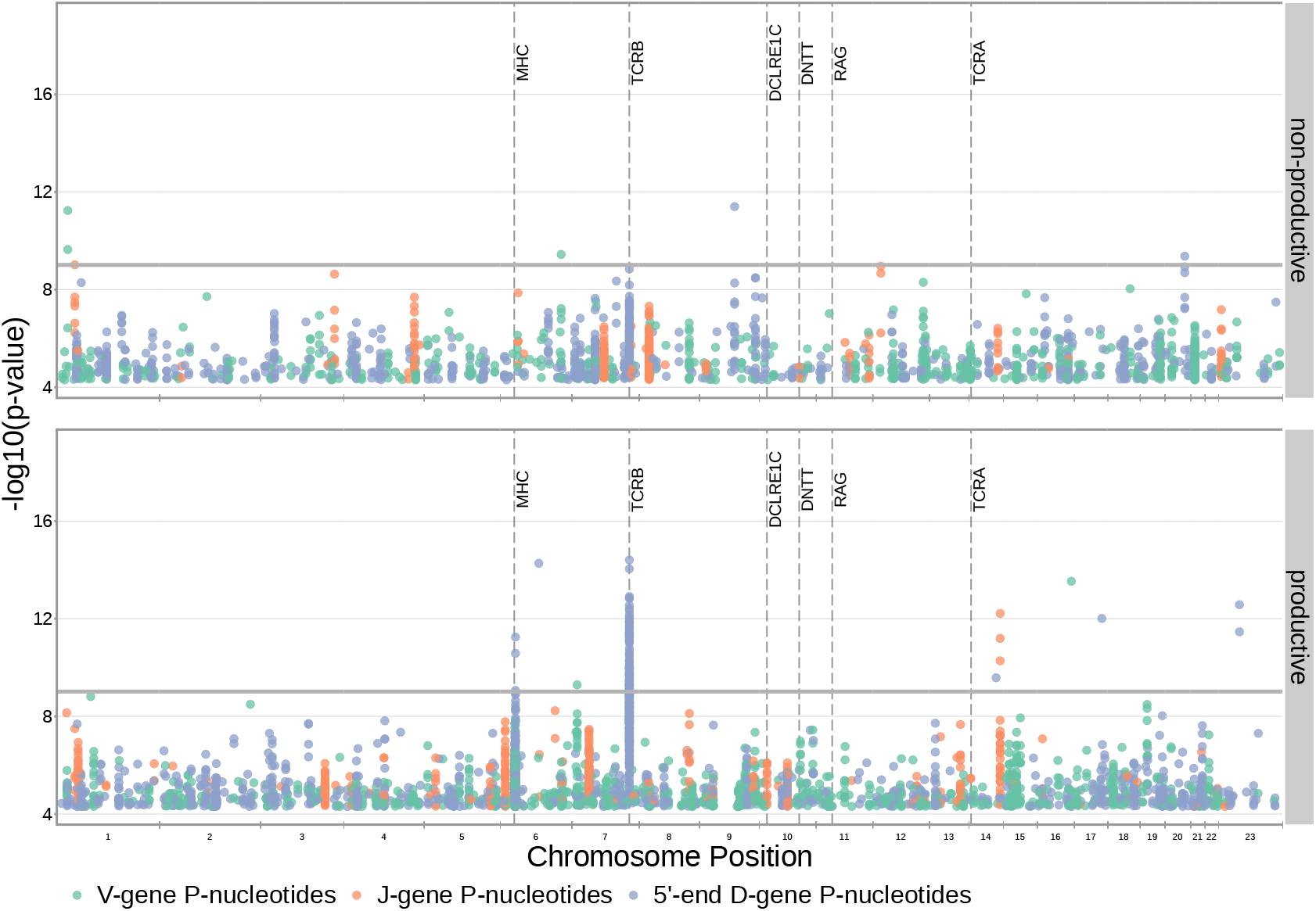
SNP associations for the number of P-nucleotides are not significant within the *DCLRE1C* locus. However, significant associations are present within the *TCRB* and MHC loci. Only SNP associations whose *P* < 5 × 10^−5^ are shown here. The gray horizontal line corresponds to a Bonferroni-corrected whole-genome P-value significance threshold of 9.68 × 10^−10^ (see Methods).

**Table 2-Figure supplement 4.**
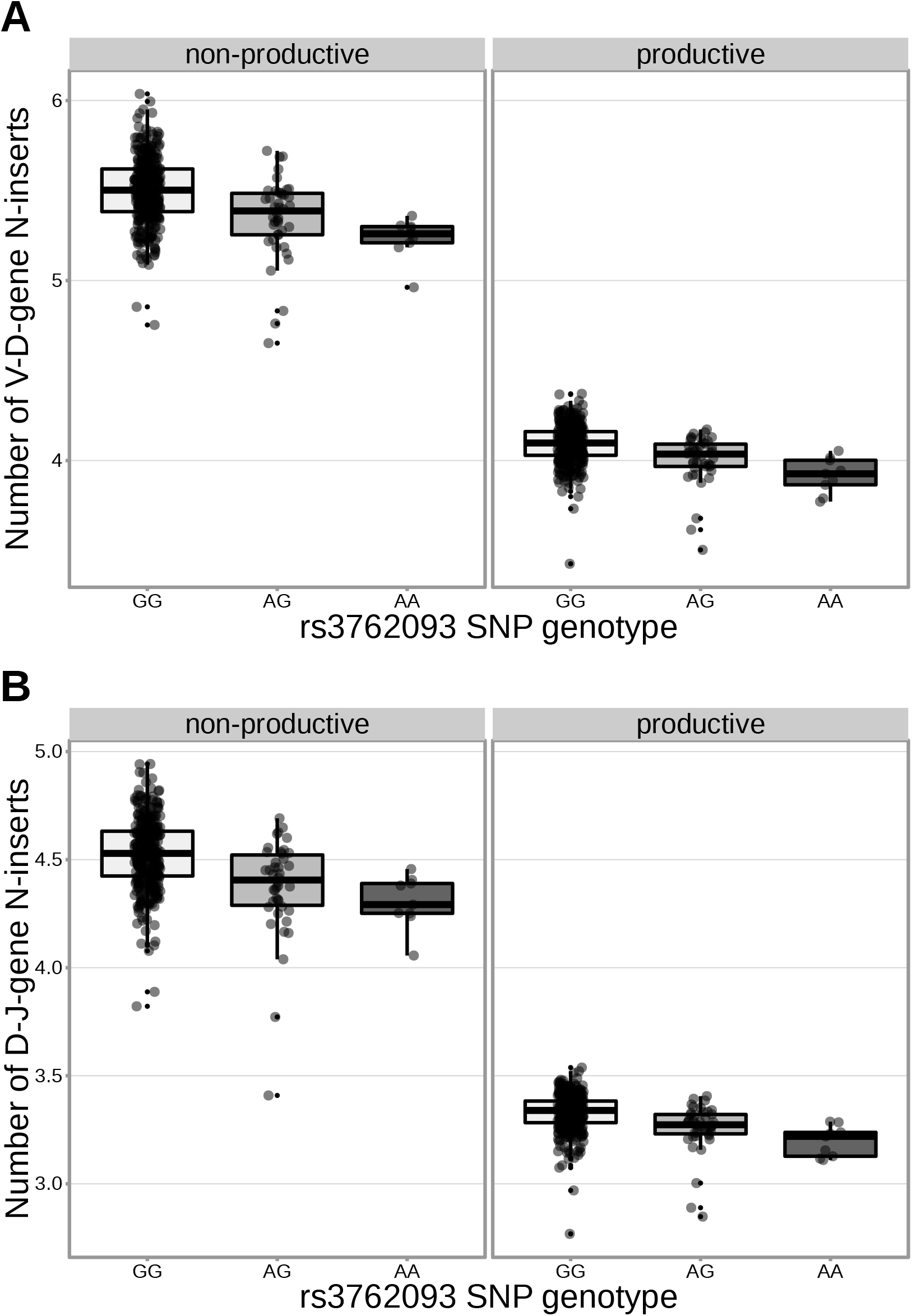
The extent of V-D and D-J N-insertion of productive and non-productive TCR*β* chains changes as a function of SNP genotype within the discovery cohort for an intronic *DNTT* SNP (rs3762093). The average number of N-insertions was calculated across all TCR*β* chains for each subject.

**Figure 4-Figure supplement 1.**
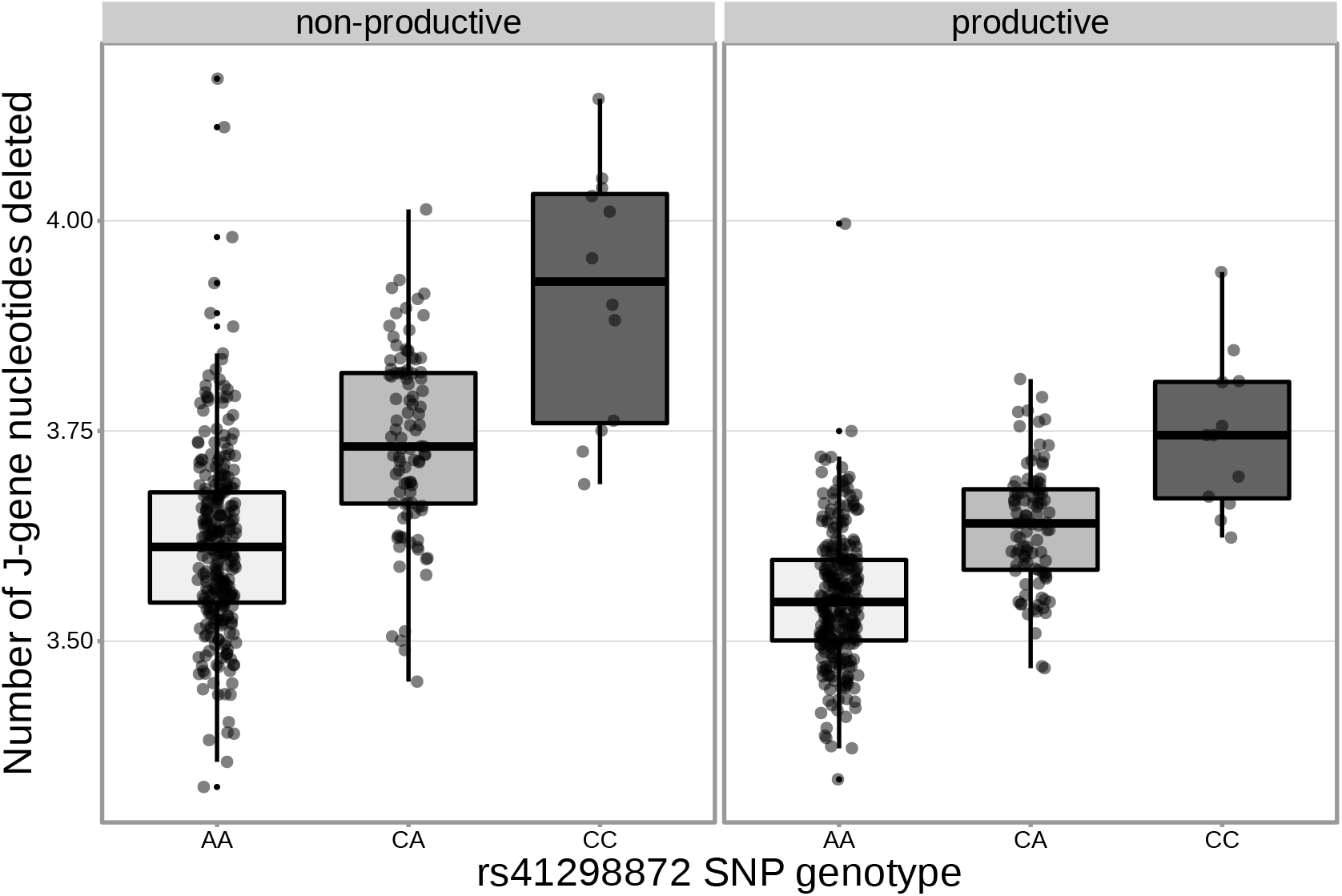
The extent of J-gene trimming changes as a function of SNP genotype for the SNP (rs41298872) most significantly associated with J-gene trimming within the *DCLRE1C* locus. Only TCRs containing *TRBJ1-1*01* (the most frequently used *TRBJ1* gene across subjects) were included when calculating the average number of J-gene nucleotides deleted for each subject.

**Table 2-Figure supplement 5.**
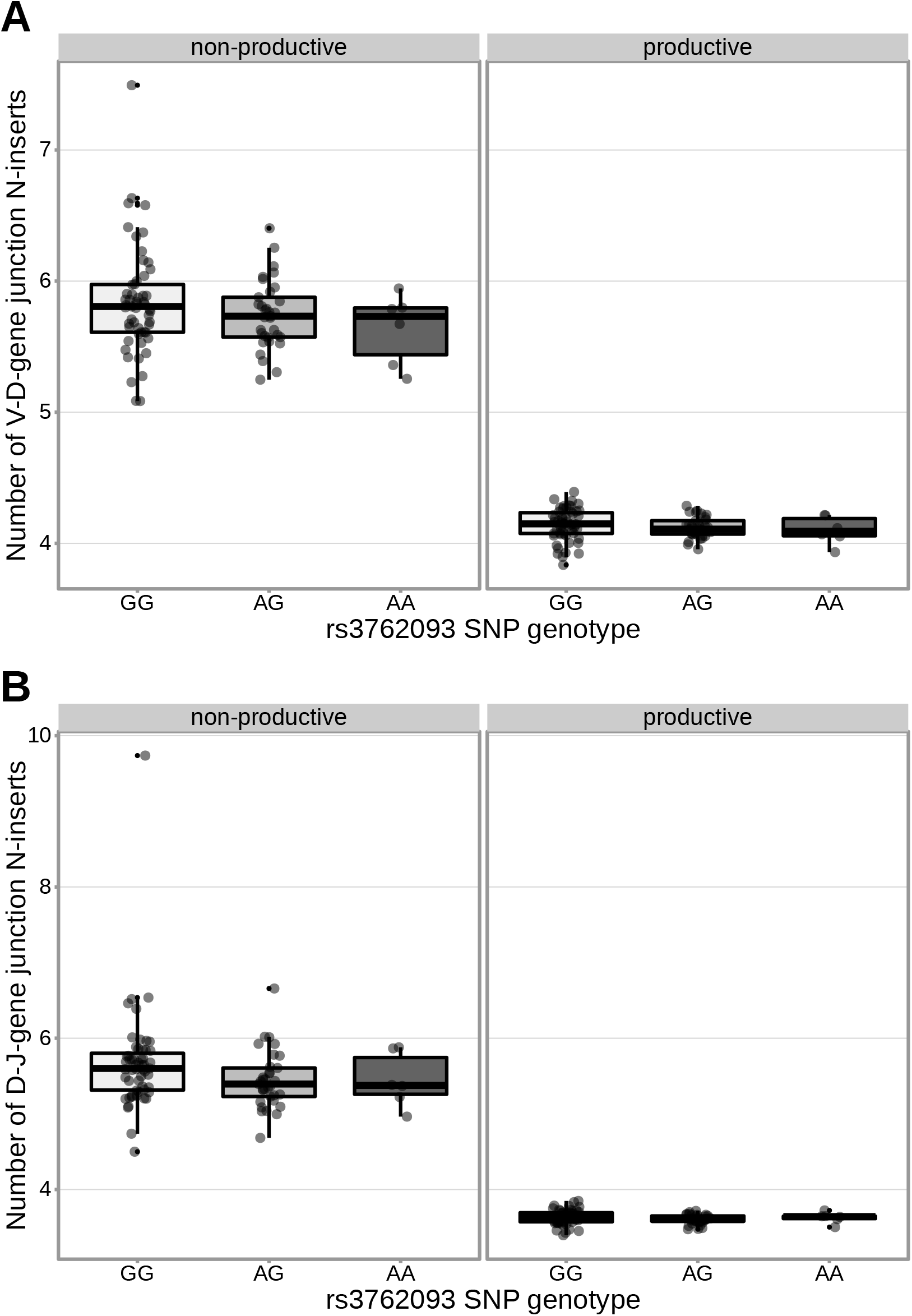
An intronic SNP (rs3762093) within the *DNTT* gene locus is not strongly associated with the number of V-D (**A**) or D-J (**B**) N-inserts within productive or non-productive TCR*β* chains in the validation cohort. However, the direction of the effect is the same as the discovery cohort for all N-insertion and productivity types. The average number of N-insertions was calculated across all TCR*β* chains for each subject.

**Figure 5-Figure supplement 1.**
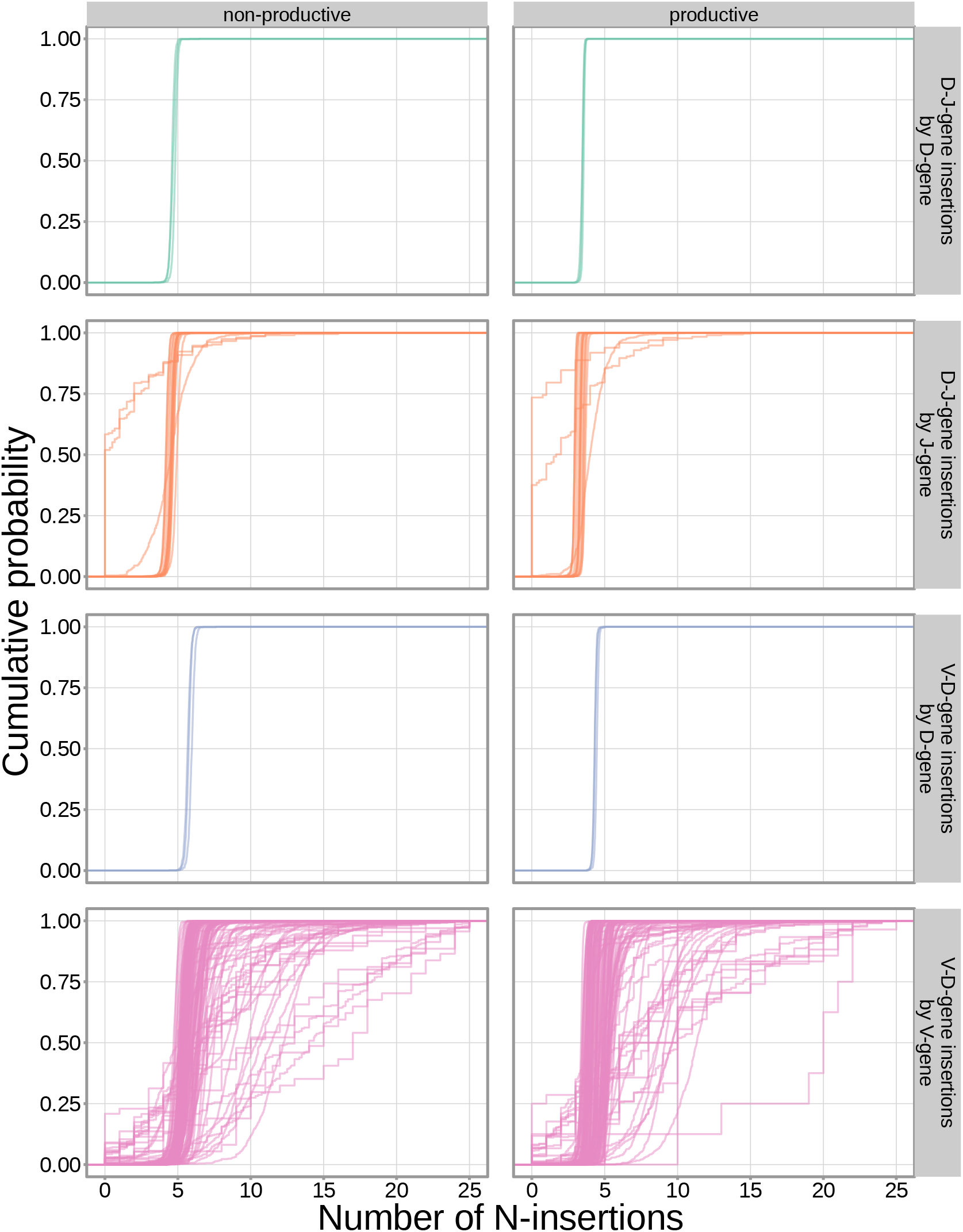
The extent of N-insertion does not vary substantially by the gene allele identity for any gene type. An empirical cumulative distribution is drawn for each gene allele type within each indicated gene type (i.e. V-gene, D-gene, J-gene).

**Figure 5-Figure supplement 2.**
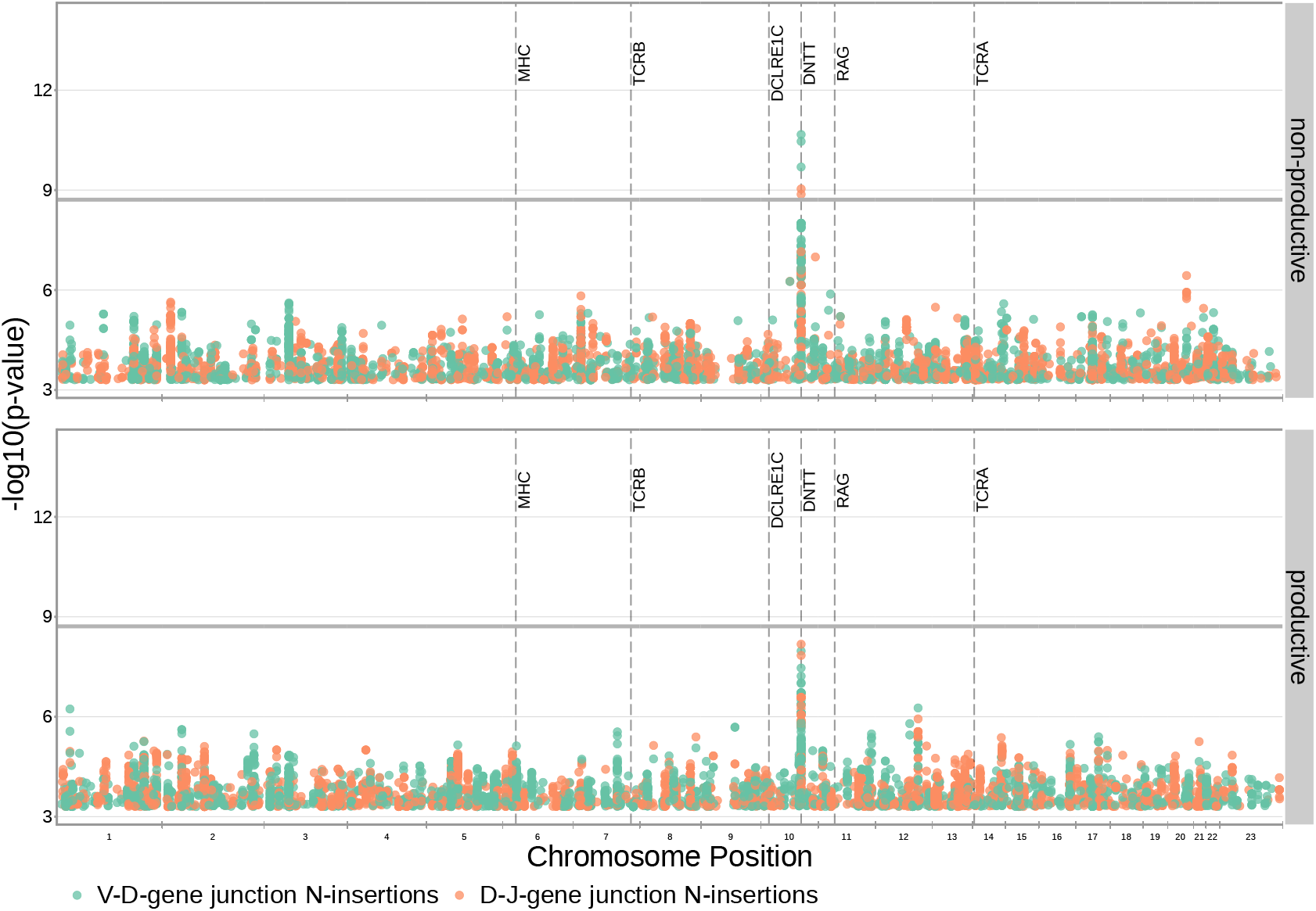
Significant associations continue to be observed within the *DNTT* locus for both V-D- and D-J-gene-junction N-insertions when restricting the analysis to TCRs which contain *TRBJ1* genes (and consequently contain *TRBD1*. Only SNP associations whose *P* < 5 × 10^−4^ are shown here. The gray horizontal line corresponds to a Bonferroni-corrected P-value significance threshold of 1.94 × 10^−9^ (calculated using whole-genome Bonferroni correction, see Methods).

**Table 2-Figure supplement 6.**
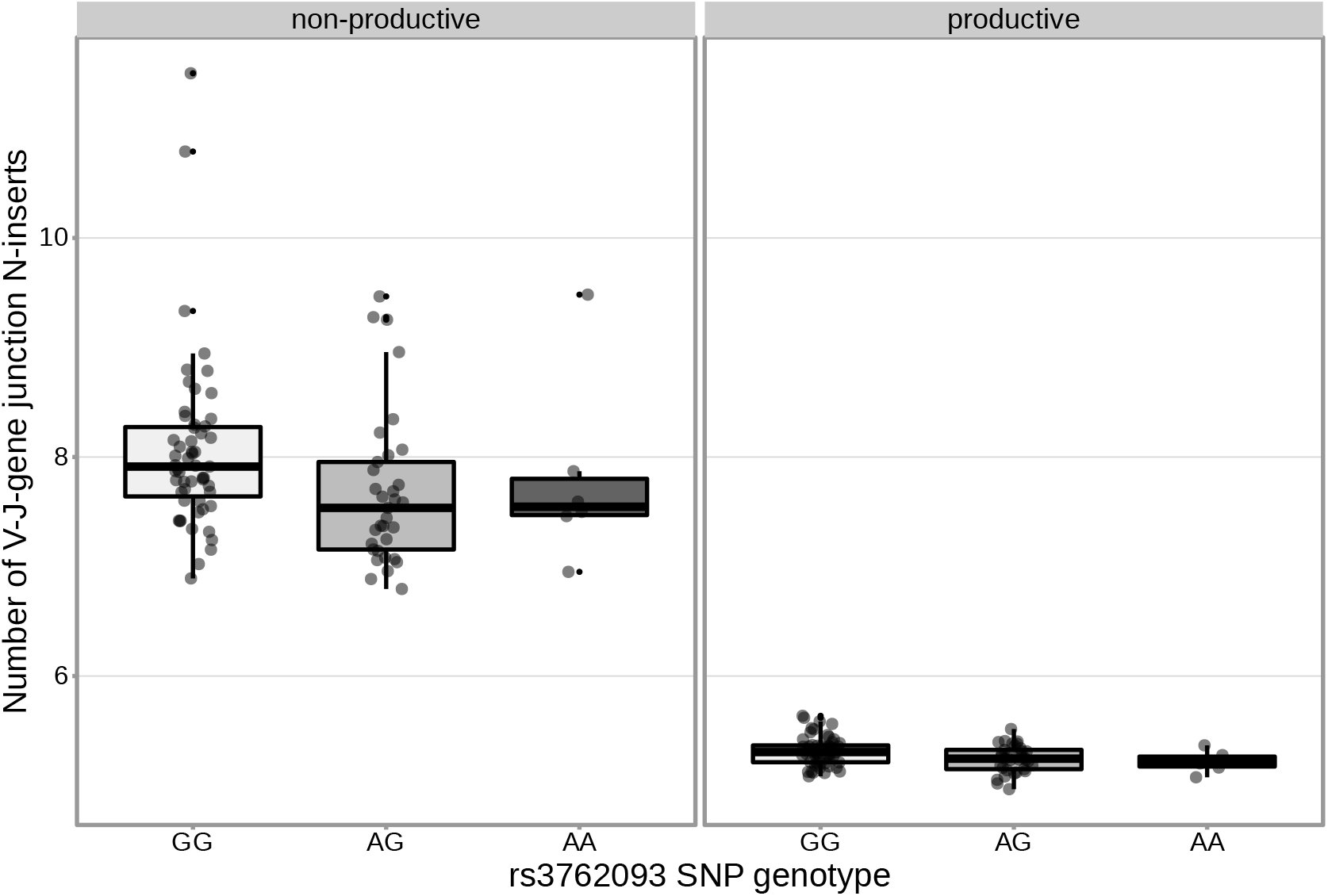
An intronic SNP (rs3762093) within the *DNTT* gene locus is significantly associated with the number of V-J N-inserts for productive TCR*α* chains in the validation cohort. This SNP is not significantly associated with the number of V-J N-inserts for non-productive TCR*α* chains in the validation cohort. The average number of N-insertions was calculated across all TCR*α* chains for each subject.

**Figure 7-Figure supplement 1.**
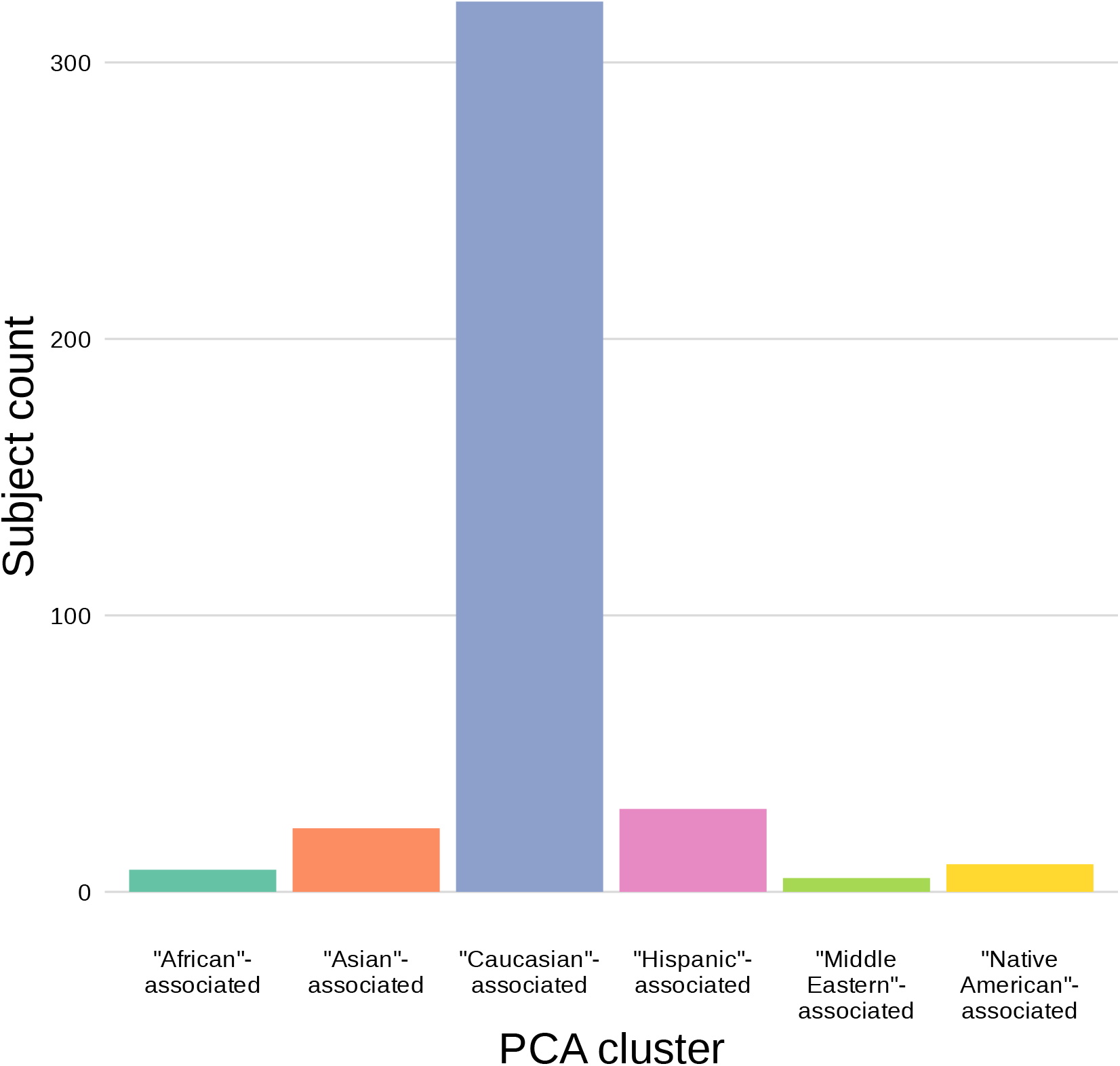
The population mean is dominated by subjects in the “Caucasian”- associated PCA-cluster. Of the 398 subjects in the sample population, 81% are in the “Caucasian”- associated PCA-cluster.

**Figure 1-source data 1.** There are 9,957 significant associations between the frequency of various V-, D-, andJ-genes and the genotype of SNPs genome-wide. The model type and Bonferroni-corrected P-value significance threshold used to identify these significant associations are described in ***Table 1***.

**Figure 1-source data 2.** Genomic inflation factor values are less than 1.03for all paired gene-frequency, productivity GWAS analyses. This suggests that we have properly controlled for population-substructure-related biases in all of the gene usage analyses. Genomic inflation factor values were calculated as described in Methods.

**Figure 2-source data 1.** Top *TCRB* SNP, gene-usage association P-value for each different V-gene, D-gene, and J-gene.

**Figure 3-source data 1.** There are 317 significant SNP associations with the extent of nucleotide trimming for various trimming types. The model type and Bonferroni-corrected P-value significance threshold used to identify these significant associations are described in ***Table 1***.

**Figure 3-source data 2.** Genomic inflation factor values are less than 1.03 for all paired nucleotide trimming, productivity GWAS analyses. This suggests that we have properly controlled for population-substructure-related biases in all of the nucleotide trimming analyses. Genomic inflation factor values were calculated as described in Methods.

**Figure 4-source data 1.** *DCLRE1C* locus SNP association P-values and locus positions.

**Figure 4-source data 2.** There are two independent SNP signals within the *DCLRE1C* locus for J-gene trimming of non-productive TCRs. A conditional analysis was performed (as described in Methods) to identify these independent signals.

**Figure 5-source data 1.** There are three significant associations between SNPs genome-wide and the number of nucleotide N-insertions. The model type and Bonferroni-corrected P-value significance threshold used to identify these significant associations are described in ***Table 1***.

**Figure 5-source data 2.** There are 232 significant associations between SNPs genome-wide and the number of nucleotide N-insertions when restricting the analysis to the extended *DNTT* locus.

**Figure 5-source data 3.** Genomic inflation factor values are less than 1.03 for all paired N-insertion, productivity GWAS analyses. This suggests that we have properly controlled for population-substructure-related biases in all of the N-insertion analyses. Genomic inflation factor values were calculated as described in Methods.

**Figure 6-source data 1.** *DNTT* locus SNP association P-values and locus positions.

**Figure 7-source data 1.** PCA-cluster and average number of N-insertions by subject.

**Figure 8-source data 1.** Minor allele frequency for N-insertion associated SNPs within the *DNTT* locus by PCA-cluster.

**Figure 9-source data 1.** Percent variance explained by each principal component.

**Figure 9-source data 2.** Scaled principal component values by subject.

## References

Conomos MP, Laurie CA, Stilp AM, Gogarten SM, McHugh CP, Nelson SC, Sofer T, Fernández-Rhodes L, Justice AE, Graff M, Young KL, Seyerle AA, Avery CL, Taylor KD, Rotter JI, Talavera GA, Daviglus ML, Wassertheil-Smoller S, Schneiderman N, Heiss G, et al. Genetic Diversity and Association Studies in US Hispanic/Latino Populations: Applications in the Hispanic Community Health Study/Study of Latinos. Am J Hum Genet. 2016 Jan; 98(1):165–184.

Conomos MP, Miller MB, Thornton TA. Robust inference of population structure for ancestry prediction and correction of stratification in the presence of relatedness. Genet Epidemiol. 2015 May; 39(4):276–293.

Corporation M, Weston S. doParallel: Foreach Parallel Adaptor for the ‘parallel’ Package; 2020, https://CRAN.R-project.org/package=doParallel, r package version 1.0.16.

Dash P, Fiore-Gartland AJ, Hertz T, Wang GC, Sharma S, Souquette A, Crawford JC, Clemens EB, Nguyen THO, Kedzierska K, La Gruta NL, Bradley P, Thomas PG. Quantifiable predictive features define epitope-specific T cell receptor repertoires. Nature. 2017 Jul; 547(7661):89–93.

Devlin B, Roeder K. Genomic control for association studies. Biometrics. 1999 Dec; 55(4):997–1004.

DeWitt WS 3rd, Smith A, Schoch G, Hansen JA, Matsen FA 4th, Bradley P. Human T cell receptor occurrence patterns encode immune history, genetic background, and receptor specificity. Elife. 2018 Aug; 7.

Dowle M, Srinivasan A. data.table: Extension of ‘data.frame’; 2021, https://CRAN.R-project.org/package=data.table, r package version 1.14.0.

Egorov ES, Merzlyak EM, Shelenkov AA, Britanova OV, Sharonov GV, Staroverov DB, Bolotin DA, Davydov AN, Barsova E, Lebedev YB, Shugay M, Chudakov DM. Quantitative profiling of immune repertoires for minor lymphocyte counts using unique molecular identifiers. J Immunol. 2015 Jun; 194(12):6155–6163.

Emerson RO, DeWitt WS, Vignali M, Gravley J, Hu JK, Osborne EJ, Desmarais C, Klinger M, Carlson CS, Hansen JA, Rieder M, Robins HS. Immunosequencing identifies signatures of cytomegalovirus exposure history and HLA-mediated effects on the T cell repertoire. Nat Genet. 2017 May; 49(5):659–665.

Feeney AJ, Victor KD, Vu K, Nadel B, Chukwuocha RU. Influence of the V(D)J recombination mechanism on the formation of the primary T and B cell repertoires. Semin Immunol. 1994 Jun; 6(3):155–163.

Fischer S, Stanke F, Tümmler B. VJ Segment Usage of TCR-Beta Repertoire in Monozygotic Cystic Fibrosis Twins. Front Immunol. 2021 Feb; 12:599133.

Freedman ML, Reich D, Penney KL, McDonald GJ, Mignault AA, Patterson N, Gabriel SB, Topol EJ, Smoller JW, Pato CN, Pato MT, Petryshen TL, Kolonel LN, Lander ES, Sklar P, Henderson B, Hirschhorn JN, Altshuler D. Assessing the impact of population stratification on genetic association studies. Nat Genet. 2004 Apr; 36(4):388–393.

Fugmann SD, Lee AI, Shockett PE, Villey IJ, Schatz DG. The RAG proteins and V(D)J recombination: complexes, ends, and transposition. Annu Rev Immunol. 2000; 18(1):495–527.

Gao K, Chen L, Zhang Y, Zhao Y, Wan Z, Wu J, Lin L, Kuang Y, Lu J, Zhang X, Tian L, Liu X, Qiu X. Germline-encoded TCR-MHC contacts promote TCR V gene bias in umbilical cord blood T cell repertoire. Front Immunol. 2019 Aug; 10:2064.

Gauss GH, Lieber MR. Mechanistic constraints on diversity in human V(D)J recombination. Mol Cell Biol. 1996 Jan; 16(1):258–269.

Gellert M. DNA double-strand breaks and hairpins in V(D)J recombination. Semin Immunol. 1994 Jun; 6(3):125–130.

Gilfillan S, Dierich A, Lemeur M, Benoist C, Mathis D. Mice lacking TdT: mature animals with an immature lymphocyte repertoire. Science. 1993 Aug; 261(5125):1175–1178.

Giudicelli V, Chaume D, Lefranc MP. IMGT/GENE-DB: a comprehensive database for human and mouse immunoglobulin and T cell receptor genes. Nucleic Acids Res. 2005 Jan; 33(Database issue):D256–61.

Gogarten SM, Bhangale T, Conomos MP, Laurie CA, McHugh CP, Painter I, Zheng X, Crosslin DR, Levine D, Lumley T, Nelson SC, Rice K, Shen J, Swarnkar R, Weir BS, Laurie CC. GWASTools: an R/Bioconductor package for quality control and analysis of genome-wide association studies. Bioinformatics. 2012; 28(24):3329–3331. doi: 10.1093/bioinformatics/bts610.

Gogarten SM, Sofer T, Chen H, Yu C, Brody JA, Thornton TA, Rice KM, Conomos MP. Genetic association testing using the GENESIS R/Bioconductor package. Bioinformatics. 2019; doi: 10.1093/bioinformatics/btz567.

Goldrath AW, Bevan MJ. Selecting and maintaining a diverse T-cell repertoire. Nature. 1999 Nov; 402(6759):255–262.

Gu J, Li S, Zhang X, Wang LC, Niewolik D, Schwarz K, Legerski RJ, Zandi E, Lieber MR. DNA-PKcs regulates a single-stranded DNA endonuclease activity of Artemis. DNA Repair (Amst). 2010 Apr; 9(4):429–437.

Jackson KJL, Gaeta B, Sewell W, Collins AM. Exonuclease activity and P nucleotide addition in the generation of the expressed immunoglobulin repertoire. BMC Immunol. 2004 Sep; 5:19.

Kallenbach S, Doyen N, Fanton d’Andon M, Rougeon F. Three lymphoid-specific factors account for all junctional diversity characteristic of somatic assembly of T-cell receptor and immunoglobulin genes. Proc Natl Acad Sci U S A. 1992 Apr; 89(7):2799–2803.

Komori T, Okada A, Stewart V, Alt FW. Lack of N regions in antigen receptor variable region genes ofTdT-deficient lymphocytes. Science. 1993 Aug; 261(5125):1171–1175.

Krishna C, Chowell D, Gönen M, Elhanati Y, Chan TA. Genetic and environmental determinants of human TCR repertoire diversity. Immun Ageing. 2020 Dec; 17(1).

Li S, Chang HH, Niewolik D, Hedrick MP, Pinkerton AB, Hassig CA, Schwarz K, Lieber MR. Evidence that the DNA endonuclease ARTEMIS also has intrinsic 5’-exonuclease activity. J Biol Chem. 2014 Mar; 289(11):7825–7834.

Lu H, Schwarz K, Lieber MR. Extent to which hairpin opening by the Artemis:DNA-PKcs complex can contribute to junctional diversity in V(D)J recombination. Nucleic Acids Res. 2007 Oct; 35(20):6917–6923.

Ma Y, Pannicke U, Schwarz K, Lieber MR. Hairpin opening and overhang processing by an Artemis/DNA-dependent protein kinase complex in nonhomologous end joining and V(D)J recombination. Cell. 2002 Mar; 108(6):781–794.

Martin PJ, Levine DM, Storer BE, Nelson SC, Dong X, Hansen JA. Recipient and donor genetic variants associated with mortality after allogeneic hematopoietic cell transplantation. Blood Adv. 2020 Jul; 4(14):3224–3233.

Mikocziova I, Greiff V, Sollid LM. Immunoglobulin germline gene variation and its impact on human disease. Genes Immun. 2021 Jun; p. 1–13.

Moshous D, Callebaut I, de Chasseval R, Corneo B, Cavazzana-Calvo M, Le Deist F, Tezcan I, Sanal O, Bertrand Y, Philippe N, Fischer A, de Villartay JP. Artemis, a novel DNA double-strand break repair/V(D)J recombination protein, is mutated in human severe combined immune deficiency. Cell. 2001 Apr; 105(2):177–186.

Murphy K, Weaver C. Janeway’s Immunobiology. Garland Science; 2016.

Murugan A, Mora T, Walczak AM, Callan CG Jr. Statistical inference of the generation probability of T-cell receptors from sequence repertoires. Proc Natl Acad Sci U S A. 2012 Oct; 109(40):16161–16166.

Nadel B, Feeney AJ. Influence of coding-end sequence on coding-end processing in V(D)J recombination. J Immunol. 1995 Nov; 155(9):4322–4329.

Nadel B, Feeney AJ. Nucleotide deletion and P addition in V(D)J recombination: a determinant role of the coding-end sequence. Mol Cell Biol. 1997 Jul; 17(7):3768–3778.

Ng S, Lopez R, Kuan G, Gresh L, Balmaseda A, Harris E, Gordon A. The timeline of influenza virus shedding in children and adults in a household transmission study of influenza in Managua, Nicaragua. Pediatr Infect Dis J. 2016;.

Oltz EM. Regulation of antigen receptor gene assembly in lymphocytes. Immunol Res. 2001; 23(2-3):121–133.

Pogorelyy MV, Minervina AA, Touzel MP, Sycheva AL, Komech EA, Kovalenko EI, Karganova GG, Egorov ES, Komkov AY, Chudakov DM, Mamedov IZ, Mora T, Walczak AM, Lebedev YB. Precise tracking of vaccineresponding T cell clones reveals convergent and personalized response in identical twins. Proc Natl Acad Sci U S A. 2018 Dec; 115(50):12704–12709.

Price AL, Zaitlen NA, Reich D, Patterson N. New approaches to population stratification in genome-wide association studies. Nat Rev Genet. 2010 Jul; 11(7):459–463.

Qi Q, Cavanagh MM, Le Saux S, NamKoong H, Kim C, Turgano E, Liu Y, Wang C, Mackey S, Swan GE, Dekker CL, Olshen RA, Boyd SD, Weyand CM, Tian L, Goronzy JJ. Diversification of the antigen-specific T cell receptor repertoire after varicella zoster vaccination. Sci Transl Med. 2016 Mar; 8(332):332ra46.

Robins HS, Srivastava SK, Campregher PV, Turtle CJ, Andriesen J, Riddell SR, Carlson CS, Warren EH. Overlap and effective size of the human CD8+ T cell receptor repertoire. Sci Transl Med. 2010 Sep; 2(47):47ra64.

Rousseeuw PJ, Van Driessen K. A Fast Algorithm for the Minimum Covariance Determinant Estimator. Technometrics. 1999 Aug; 41(3):212–223.

Rubelt F, Bolen CR, McGuire HM, Vander Heiden JA, Gadala-Maria D, Levin M, Euskirchen GM, Mamedov MR, Swan GE, Dekker CL, Cowell LG, Kleinstein SH, Davis MM, Individual heritable differences result in unique cell lymphocyte receptor repertoires of naïve and antigen-experienced cells; 2016.

Schatz DG, Swanson PC. V(D)J recombination: mechanisms of initiation. Annu Rev Genet. 2011 Aug; 45(1):167–202.

Sharon E, Sibener LV, Battle A, Fraser HB, Garcia KC, Pritchard JK. Genetic variation in MHC proteins is associated with T cell receptor expression biases. Nat Genet. 2016 Sep; 48(9):995–1002.

Shugay M, Britanova OV, Merzlyak EM, Turchaninova MA, Mamedov IZ, Tuganbaev TR, Bolotin DA, Staroverov DB, Putintseva EV, Plevova K, Linnemann C, Shagin D, Pospisilova S, Lukyanov S, Schumacher TN, Chudakov DM. Towards error-free profiling of immune repertoires. Nat Methods. 2014 Jun; 11(6):653–655.

Srivastava SK, Robins HS. Palindromic nucleotide analysis in human T cell receptor rearrangements. PLoS One. 2012 Dec; 7(12):e52250.

Tanno H, Gould TM, McDaniel JR, Cao W, Tanno Y, Durrett RE, Park D, Cate SJ, Hildebrand WH, Dekker CL, Tian L, Weyand CM, Georgiou G, Goronzy JJ. Determinants governing T cell receptor *α*/*β*-chain pairing in repertoire formation of identical twins. Proc Natl Acad Sci U S A. 2020 Jan; 117(1):532–540.

Thomas PG, Crawford JC. Selected before selection: A case for inherent antigen bias in the T cell receptor repertoire. Curr Opin Syst Biol. 2019 Dec; 18:36–43.

Weigert M, Gatmaitan L, Loh E, Schilling J, Hood L. Rearrangement of genetic information may produce immunoglobulin diversity. Nature. 1978; 276(5690):785–790.

Wickham H, Averick M, Bryan J, Chang W, McGowan LD, François R, Grolemund G, Hayes A, Henry L, Hester J, Kuhn M, Pedersen TL, Miller E, Bache SM, Müller K, Ooms J, Robinson D, Seidel DP, Spinu V, Takahashi K, et al. Welcome to the tidyverse. Journal of Open Source Software. 2019; 4(43):1686. doi: 10.21105/joss.01686.

Wilke CO. cowplot: Streamlined Plot Theme and Plot Annotations for ‘ggplot2’; 2020, https://CRAN.R-project.org/package=cowplot, r package version 1.1.1.

Witzgall R, O’Leary E, Leaf A, Onaldi D, Bonventre JV. The Krüppel-associated box-A (KRAB-A) domain of zinc finger proteins mediates transcriptional repression. Proc Natl Acad Sci U S A. 1994 May; 91(10):4514–4518.

Woodsworth DJ, Castellarin M, Holt RA. Sequence analysis of T-cell repertoires in health and disease. Genome Med. 2013 Oct; 5(10):98.

Zhao B, Rothenberg E, Ramsden DA, Lieber MR. The molecular basis and disease relevance of non-homologous DNA end joining. Nat Rev Mol Cell Biol. 2020 Dec; 21(12):765–781.

Zheng X, Levine D, Shen J, Gogarten S, Laurie C, Weir B. A High-performance Computing Toolset for Relatedness and Principal Component Analysis of SNP Data. Bioinformatics. 2012; 28(24):3326–3328. doi: 10.1093/bioinformatics/bts606.

Zvyagin IV, Pogorelyy MV, Ivanova ME, Komech EA, Shugay M, Bolotin DA, Shelenkov AA, Kurnosov AA, Staroverov DB, Chudakov DM, Lebedev YB, Mamedov IZ. Distinctive properties of identical twins’ TCR repertoires revealed by high-throughput sequencing. Proc Natl Acad Sci U S A. 2014 Apr; 111(16):5980–5985.

